# Single-cell multiomics reveals epigenetic rewiring of splenic memory B cells in murine malaria reinfection

**DOI:** 10.64898/2025.12.10.693423

**Authors:** Montserrat Coronado, África Vincelle-Nieto, Isabel G. Azcárate, Susana Pérez-Benavente, Antonio Puyet, Amalia Díez, José M. Bautista, Armando Reyes-Palomares

**Author notes:** Corresponding Authors: Armando Reyes-Palomares, José M. Bautista.

## Abstract

Malaria induces slow, gradually acquired, non-sterilizing immunity whose cellular and regulatory underpinnings remain incompletely understood. Here, we combine a sequential *Plasmodium yoelii* 17XNL infection model in BALB/c mice, in which primary parasitemia resolves spontaneously and confers robust protection upon homologous reinfection, with single-cell RNA and chromatin accessibility profiling to dissect how primary infection and recall reshape splenic immunity, with a focus on B cells. We generate a multiomic atlas of >50,000 splenic mononuclear cells, resolving thirteen major immune lineages and 48 subpopulations, and show that B cells dominate the response and diversify into naïve/mature, germinal center, memory, and plasmablast compartments. Trajectory analysis reveals distinct differentiation paths towards germinal center, memory, and mature B cells, and uncovers infection-dependent shifts in transcription factor activity, cis-regulatory element usage, and gene regulatory networks. Reinfection is associated with a shift in memory B-cell composition and transcriptional programs towards extrafollicular-like, IgM⁻ conventional memory B cells together with epigenetic modules linked to rapid antibody production. Together, these data provide a systems-level view of B cell plasticity in experimental malaria and provides a mechanistic framework from a highly protective *P. yoelii* reinfection model with implications for understanding non-sterilizing immunity in endemic settings.

## Introduction

Malaria remains one of the most devastating infectious diseases worldwide, causing hundreds of thousands of deaths each year, predominantly in children under five in endemic regions. The disease is caused by protozoan parasites of the genus *Plasmodium*, which undergo an obligatory blood stage where they infect and replicate within red blood cells. Unlike many acute viral or bacterial infections, naturally acquired immunity to malaria in endemic populations is slow to develop^1^, requires repeated parasite exposure, and is typically non-sterilizing^2^, reducing disease severity and parasitemia without fully preventing reinfection^3,4^.

This atypical pattern of immunity, which only develops after multiple infections and still fails to completely prevent new episodes, suggests that malaria relies on unconventional mechanisms of immune memory^2^. It also raises fundamental questions about how repeated blood-stage infections progressively reshape lymphocyte differentiation programs and fine-tune their effector functions^3,5^. Although clinical and experimental studies have described aspects of B and T cell responses over the course of infection, we still lack a mechanistic, systems-level view of how primary and recall responses are encoded in the immune system during malaria.

Spleen is a central site for blood-stage parasite clearance and for the initiation and maintenance of anti-malarial humoral immunity^6–10^. Splenic architecture and cellular composition are profoundly remodeled during *Plasmodium* infection, with de novo germinal center (GC) formation and expansion within the white pulp, distortion or loss of marginal-zone and T cell compartments that varies with host and parasite strain, and marked splenomegaly as a consistent feature in both human and experimental malaria^6,7,9,11^. In severe *Plasmodium* infections or in splenectomized hosts, parasite control is impaired and mortality is markedly increased, underscoring the importance of splenic responses for survival. Murine models based on non-lethal strains such as *P. yoelii* 17XNL (Py17XNL) recapitulate key features of human malaria, including self-resolving primary parasitemia and strong protection against subsequent challenge^12,13^. These models provide a tractable system to dissect how primary and recall responses are encoded within splenic lymphocyte compartments^14,15^.

B cells are central effectors of anti-malarial immunity because they generate antibodies that recognize parasite antigens on infected erythrocytes and circulating merozoites. These antibodies opsonize infected red blood cells for phagocytic clearance in the spleen and block merozoite invasion of new erythrocytes, thereby limiting parasite replication and peak parasitemia. In addition, memory B cells established after primary infection enable a faster and higher-affinity antibody response upon reinfection, further enhancing parasite control^16^. However, B cells do not behave as a homogeneous population since they encompass naïve and mature follicular (FOB) and marginal zone cells (MZB), GC B cells, plasmablasts and plasma cells, and multiple memory B cell subsets. Memory B cells themselves are functionally and developmentally heterogeneous, arising through GC-dependent and extrafollicular pathways and differing in isotype (IgM⁺ versus class-switched IgM⁻), degree of affinity maturation, and recall potential^17–20^. Recent work in human malaria and murine *Plasmodium* models has delineated classical memory B cells (cMBCs) and atypical/age-associated B (ABCs) cells that expand during chronic infection and autoimmunity^21–24^. In *P. yoelii* infection, IgM⁺ and class-switched IgM⁻ memory B cells with extrafollicular-like phenotypes mount rapid recall responses and contribute to protection upon rechallenge^15,25,26^. Yet, how repeated *Plasmodium* exposures reprogram the transcriptional and epigenetic landscape of splenic B cell subsets, and how gene regulatory networks (GRNs) are rewired between primary and secondary responses, remains poorly understood^27,28^.

The recent advent of single-cell multiomic technologies provides an opportunity to address these questions by simultaneously measuring gene expression and chromatin accessibility in the same cells. Single-cell RNA sequencing (scRNA-seq) has been used to map immune cell states during acute malaria^27^, and single-cell ATAC sequencing (scATAC-seq) has recently been applied in experimental malaria to map chromatin accessibility and immune cell states, particularly within memory B cell compartments^15^. However, existing studies have largely focused on acute infection or on selected T or B cell subsets, and a comprehensive multiomic atlas of splenic mononuclear cells across naturally resolved primary infection and reinfection is lacking. Moreover, most GRN inference approaches aggregate all cells in a dataset, which can obscure context-specific regulatory circuits that operate along particular differentiation trajectories or in defined infection states.

Here, we combine a sequential Py17XNL infection model in BALB/c mice in which primary parasitemia resolves spontaneously and confers robust protection upon homologous reinfection, and joint scRNA-seq and scATAC-seq profiling to dissect how primary and secondary blood-stage infections remodel splenic immunity. We perform multiomic single-cell profiling of >50,000 splenic mononuclear cells across control, primary infection, and early and late reinfection, integrate transcriptomic and chromatin accessibility data to build a high-resolution atlas comprising thirteen major immune lineages and 48 subpopulations, and then focus on the dominant B cell compartment. Using trajectory analysis on chromatin accessibility, we reconstruct differentiation paths toward memory B cells, map infection-dependent changes in transcription factor (TF) activity, *cis*-regulatory element (CRE)-gene coupling, and GRNs along these lineages. By comparing primary infection and reinfection, we uncover transcriptional and epigenetic signatures of GC-derived versus extrafollicular memory, identify key TFs and regulatory modules that are differentially engaged between primary and recall responses, and reveal how repeated *Plasmodium* exposure reshapes the regulatory architecture of splenic B cell immunity. Together, these findings provide a systems-level view of B cell diversification and memory formation in an experimental model of highly protective malaria reinfection and how repeated infections may sustain non-sterilizing immunity in chronic and recurrent settings.

## Results

### Dynamics of *P. yoelii* 17XNL infection

To investigate how immune memory to *Plasmodium* reinfection is acquired, we performed successive experimental infections in female BALB/c mice using the non-lethal strain Py17XNL (Figure 1A). Eight mice were distributed into four experimental conditions: control (Ctrl), primary infection (Prim), early reinfection (EReinf), and late reinfection (LReinf). Throughout the experiment, we monitored parasitemia and body weight, thus allowing us to evaluate the physiological response of the animals during each infection episode (Figure 1B). In infected mice, parasitemia became microscopically detectable from 3 days post infection (dpi), increased progressively, and reached a peak around 13 dpi. Parasite levels then declined until they became to undetectable levels by approximately 21 dpi, indicating complete clearance of the primary infection in agreement with previous reports using the same strain in BALB/c mice. After resolution of primary infection cycle, a subset of mice was reinoculated at 34 dpi to assess the magnitude of the acquired immune response upon re-exposure. None of the reinfected mice developed microscopically detectable parasitemia (Figure 1B, top), demonstrating that the immune response generated during the primary infection is sufficient to prevent detectable parasite expansion upon secondary challenge under these conditions.

**Figure 1.**
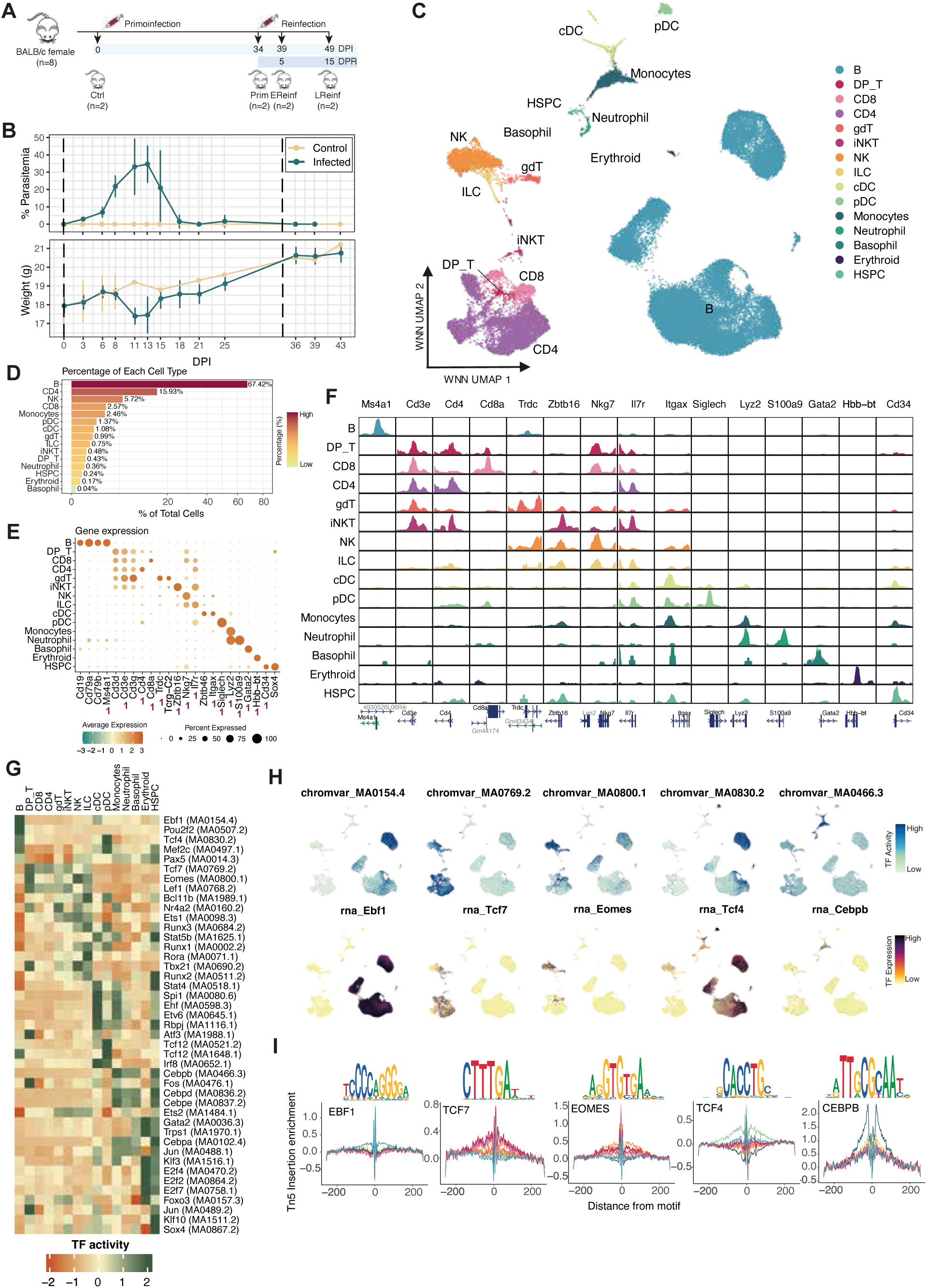
Atlas of splenic mononuclear cells in mouse. (A) Scheme of the experimental design for sequential Py17XNL infection in mice. Eight female BALB/c mice were inoculated with 1 × 10^6^ iRBCs with the non-lethal Py17XNL strain, undergoing different numbers of infection episodes. Primary infection was performed on day 0 and reinfection on day 34 post infection (dpi). According to the sacrifice time point, mice were assigned to four groups: uninfected controls, primary infection (34 dpi), early reinfection (39 dpi, i.e., 5 days post reinfection, dpr), and late reinfection (49 dpi, 15 dpr). (B) In vivo dynamics of Py17XNL infection in BALB/c mice referred to mean parasitemia (top) and body weight (bottom) of mice from the start of infection (0 dpi) to the end of the experiment. Two groups of mice are shown: infected animals and non-infected controls. Vertical dashed lines indicate the days on which primary infection (0 dpi) and reinfection (34 dpi) were performed. (C) Multimodal integration represented as WNN UMAP of the 57,842 splenic cells. Each dot corresponds to a single cell. Color indicates the major cell type. (D) Percentage of each major population relative to the total number of cells in the atlas, ordered from most to least abundant. The color gradient indicates the percentage of cells. (E) Mean expression of marker genes in each cell population. Dot size indicates the proportion of cells of that type expressing the gene, and the color gradient indicates standardized mean expression. Genes marked with an asterisk are those displayed in panel F. (F) Chromatin accessibility profiles for marker genes for each cell type. The gene name is shown at the top, and the promoter position and gene orientation are indicated at the bottom. Peak color denotes the corresponding cell type. (G-I) Identification of transcription factors associated with cell identity. (G) Heatmap showing the activity of the top 5 TFs specific to each cell type. The color gradient indicates the mean activity value in each cell type. (H) UMAP plots showing TF activity (top) and TF expression (bottom) for TFs specific to the different cell types. (I) TF footprinting analysis for selected TFs, split by cell type. Color indicates the corresponding cell type.

Body weight, increased progressively over time in both control and infected animals, consistent with normal growth (Figure 1B, bottom). In infected mice, this trend was transiently interrupted by a reduction in body weight between 8 and 11 dpi that coincided with peak of parasitemia, highlighting the marked pathophysiological impact of the acute phase.

We also monitored spleen weight as a proxy for immune activation and infection-induced inflammation. During acute infection, spleen weight in infected mice increased by more than 400% at 8 dpi, consistent with splenomegaly as a hallmark of blood-stage malaria infection and a reflection of intense lymphoid and myeloid activation (Figure S1A). Following parasite clearance and during reinfection, spleen weight gradually returned towards values observed in control mice, in line with resolution of the inflammatory response.

### Generation of an atlas of splenic mononuclear cells associated with infection and reinfection in mice

This experimental design enabled an in-depth analysis of the cellular and molecular processes occurring in the spleen during the development of immunological memory to malaria. To characterize the heterogeneity of splenic mononuclear cells and to link molecular phenotypes to specific cell subpopulations across different parasite exposures, we performed single-cell assays simultaneously profiling gene expression and chromatin accessibility using the 10x Genomics Multiome platform^29,30^. After preprocessing and quality control, we obtained 57,842 cells from the eight mice, enabling quantification of 17,584 genes and identification of 156,396 accessible chromatin regions.

To identify shared cell types across samples and control for batch-related technical variability, we first integrated gene expression and chromatin accessibility data separately for each sample (Figure S1B). We then performed multimodal integration of both omics by constructing modality-specific k-nearest neighbor graphs and combining them using the WNN framework^31^. This multimodal integration embedded each cell into a common space defined jointly by its transcriptomic and chromatin accessibility profiles (Figure 1C and Figure S1C). The resulting UMAPs showed a homogeneous distribution of cells across infection groups, while still revealing marked differences when comparing uninfected control cells with infected ones (Figure S1D).

Using the integrated multiomic data, we defined cell identities and resolved 48 distinct cellular subpopulations (Figure S1E). These subpopulations were grouped into 15 major immune cell types and subtypes (Figure 1C) and manually annotated based on expression of canonical marker genes (Figure 1E). B cells, the predominant population (67% of total cells; Figure 1D), were identified by expression of *Cd19*, *Cd79*, and *Ms4a1*. T cells expressed *Cd3d/e/g* and were further subdivided into CD4⁺ T cells (*Cd4*), CD8⁺ T cells (*Cd8a*), and a small fraction of double-positive T cells (DP_T) with reduced *Cd4* and *Cd8a* expression. Additional lymphoid subsets included γδ T cells (Tγδ; *Trdc, Tcrg-C2*), invariant NKT cells (*Zbtb16*), NK cells, and ILCs, the latter two marked by *Nkg7*.

Although myeloid cells represented a minority (5.7%), we identified monocytes (*Lyz2*), conventional dendritic cells (cDCs; *Itgax, Zbtb46*), plasmacytoid dendritic cells (pDCs; *Siglech*), neutrophils (*S100a9*), basophils (*Gata2*), erythroid cells (*Hbb-bt*), and hematopoietic stem and progenitor cells (HSPCs; *Cd34, Sox4*). Chromatin accessibility patterns showed high concordance with transcriptomic data, with increased accessibility at promoters of marker genes in the corresponding cell types, confirming at the epigenetic level the identities inferred from scRNA-seq (Figure 1F).

Complementary differential analyses of gene expression and chromatin accessibility across major cell types identified sets of genes and regions acting as population-specific markers, thereby defining characteristic cellular profiles (Figure S2A). In total, we detected 8,777 differentially expressed genes (DEGs; log2FC > 0, FDR < 0.05) and 2,572 differentially accessible regions (DARs; log2FC > 0, FDR < 0.05). Because these markers were obtained from all samples combined, we computed module scores, defined as the average expression or chromatin accessibility of each cell-type-specific DEG or DAR set in individual cells, to verify that they reflected cell identity rather than infection status. Module scores showed strong and consistent enrichment within their corresponding cell types, irrespective of experimental condition (Figure S2B), underscoring the robustness and physiological relevance of the identified markers.

We next evaluated correlations between samples using pseudobulk expression and chromatin accessibility profiles, generated by aggregating all cells from each sample. This approach provided a robust view of sample-level clustering. In each omics, we observed high overall correlation among samples but a clear separation by infection group: one cluster comprised primary infection samples, whereas another grouped control and late reinfection samples (Figure S2C). This pattern indicated that infection status, and particularly primary infection, exerts a major influence on the global organization of molecular profiles. In contrast, the two early reinfection samples showed divergent behavior: one (EReinf2) clustered with primary infection, whereas the other (EReinf1) grouped with control and late reinfection samples. Although both early reinfection samples were retained in the splenic atlas to preserve continuity along the infection course, their intermediate and variable characteristics led us to exclude them from downstream heterogeneity and differential analyses. Unless otherwise specified, the term “reinfection” hereafter refers to late reinfection samples.

### Chromatin-based inference of TF activity and cell-type annotations

Chromatin accessibility data can be leveraged to infer potential TF binding by analyzing motifs present within accessible regions. We calculated the mean accessibility signal across all binding sites associated with each TF, as an estimate of its activity, and compared these values across cell types, analogous to marker identification but focused on TF specificity. To infer TF activity per cell and quantify its cell-type specificity, we used chromVAR^32^. Differential analysis combining AUC values for gene expression and TF activity identified TFs with the highest specificity for each population.

Several TFs with well-established regulatory roles in immune lineages emerged from this analysis (Figure 1G). For example, EBF1 and PAX5 showed strong specificity in B cells, whereas TCF7 activity was distinctive of T cells. In NK and Tγδ cells, EOMES-associated activity stood out as particularly specific. In pDCs, TCF4 displayed a characteristic activity profile, with signal also detectable in a subset of B cells. CEBPB was identified as specifically active in monocytes. In many cases, strong TF activity specificity was mirrored by higher expression of the corresponding TF, reinforcing its likely regulatory role (Figure 1H).

To further support these inferences, we performed TF footprinting analysis to assess direct TF occupancy patterns in the cell types with highest inferred activity (Figure 1I). In all cases, we observed characteristic protection profiles against Tn5 insertion around binding motifs, indicating direct TF-DNA interaction at these sites. The depth and symmetry of these footprints supported the specificity and robustness of the signal, whereas differences in footprint width suggested variation in binding affinity or occupancy frequency among motifs and cell types. Notably, EBF1 and EOMES motifs showed more pronounced footprints, consistent with stable binding in B cells and NK/Tγδ cells, respectively, whereas TCF4 footprints were more attenuated, likely reflecting occupancy restricted to specific pDC subpopulations. Because scATAC-seq data are sparse footprints were computed from aggregated cells of the same type and therefore reflect average occupancy rather than single-cell TF activity. The concordance between occupancy profiles, chromVAR-inferred activity, and TF gene expression supports the existence of cell-type-specific regulatory programs aligned with immune cell identity.

To validate the accuracy of our manual annotations, we compared them to automatic annotations obtained by mapping our atlas transcriptomes onto the murine immune cell reference ImmGen^33^. Similarity between manually and automatically annotated cell groups was quantified using the Jaccard index, defined as the fraction of shared cells relative to the total number of cells in both sets. The Jaccard index was very high for major cell types, indicating strong agreement between the two independent annotation strategies (Figure S2D). However, some cell subtypes, such as Tγδ cells, iNKT cells, and ILCs, showed low similarity (Figure S2D). When the analysis was stratified by experimental condition, the group with the largest discrepancies between annotation systems was the primary Py17XNL infection group (Figure S2E). In contrast, cell subtypes in control samples showed higher similarity indices, suggesting that the most pronounced differences are associated with infection-induced immune remodeling (Figure S2E). Manual clustering and annotation also allowed us to distinguish an erythroid population that was incorrectly labeled as hematopoietic stem cells by automatic annotation.

Once the major cell types in our atlas were established, we evaluated how cellular heterogeneity changed across experimental conditions (control, primary infection, and reinfection). We compared the relative fraction of each population with respect to the total number of cells per sample (Figure S2F). This analysis revealed an increased proportion of B cells in infected samples compared with controls, particularly in primary infection, where B cells were ∼10% more abundant than in control mice. Tγδ cells showed a similar trend to B cells in infected groups, although they remained rare overall, representing only 0.99% of all cells (Figure 1D). For T-cell populations (DP_T, CD8, and CD4), we observed a progressive increase in their proportions following the temporal course of infection, from control to reinfection. In contrast, NK cells, iNKT cells, and myeloid populations were approximately twice as abundant in control samples as in infected samples.

### Characterization of cis-regulatory elements remodeling and co-accessible chromatin domains

A key advantage of joint multiomic profiling in the same cells is the ability to detect regulatory interactions between CREs and their target genes^34^. To infer such links, we computed correlations between chromatin accessibility at individual peaks and the expression of genes whose transcription start sites (TSSs) were located within 500 kb. This analysis yielded 26,871 putative activating CRE-gene links (R > 0), involving 16,119 unique CREs and 5,107 distinct genes, in which correlated peaks are likely to correspond to functional CREs (Figure 2A). These correlations, derived from matched chromatin accessibility and gene expression, aligned with cell-type-specific transcriptional programs, highlighting their potential relevance for regulating physiological processes in each lineage. Across all interactions, the median number of positively correlated CREs per gene was three, whereas most CREs associated with a single gene (median = 1) (Figure 2B). Some genes, however, were associated with an exceptionally high number of CREs. Notably, *Cd28*, a T cell-expressed co-stimulatory receptor essential for TCR-mediated activation^35^, showed positive correlations with 48 CREs (Figure 2C).

**Figure 2.**
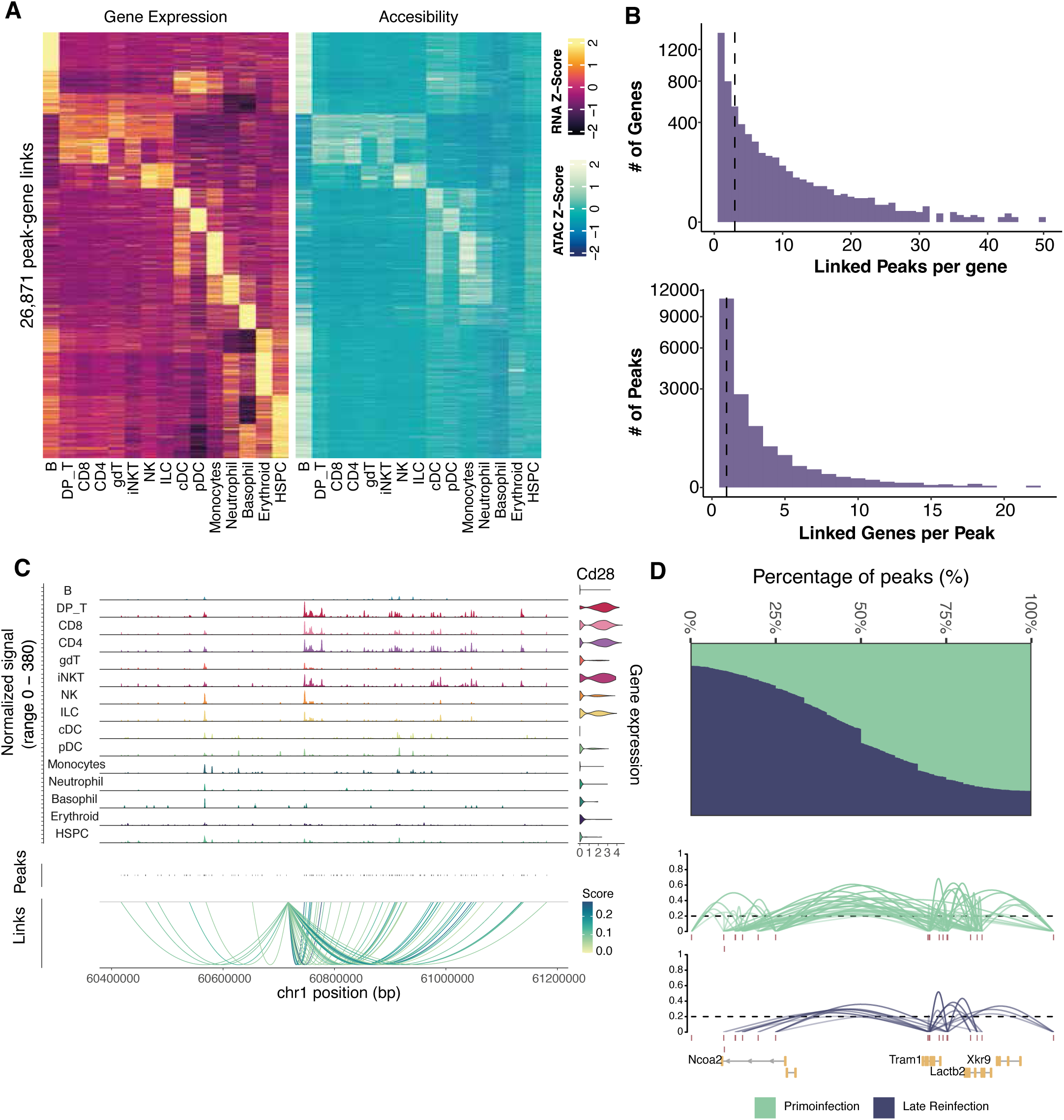
Characterization of CREs in splenic mononuclear cells. (A) Heatmaps showing gene expression (left) and chromatin accessibility (right) for peak-gene links. Each column corresponds to a pseudobulk profile for a given cell type, and each row represents a gene (left) and the accessibility of its associated CREs (right). The color gradient indicates standardized expression and accessibility values. (B) Histograms showing the distribution of the number of CREs significantly associated with each gene (top) and the number of genes associated with each peak (bottom). The dashed line indicates the median of each distribution. (C) Accessibility of CREs associated with *Cd28* across major cell types (left). On the right, a violin plot shows *Cd28* expression in each cell type. The bottom panel depicts the correlation between each regulatory peak and *Cd28* expression. (D) Identification of chromatin co-accessibility domains. (D, top) Barplot showing the distribution of the percentage of peaks with stronger co-accessibility in primary infection samples (green) or reinfection samples (blue) for each CCAN. (D, bottom) Plots of peak-peak connections within representative CCANs inferred from primary infection samples (top, green) and reinfection samples (bottom, blue). The dashed line indicates the minimum co-accessibility threshold between peaks. Co-accessibility values are encoded by line opacity.

Having established CRE-gene interactions, we next asked whether higher-order chromatin organization was modulated by infection. We used Cicero^36^ to infer domains of co-accessibility between peaks, based on the premise that peaks with coordinated accessibility across the same cell populations are likely to be functionally connected, such as an enhancer and a gene promoter. Using all cells in the atlas, we identified 2,382 cis-co-accessibility networks (CCANs) comprising groups of peaks whose accessibility covaries across cells (Figure 2D). Most genes were associated with a limited number of peaks within a CCAN, but a subset of loci displayed more extensive regulatory networks, consistent with infection-driven reorganization of chromatin accessibility profiles (Figure 2D, top). For instance, a CCAN controlling *Tram1*, *Xkr9*, *Lactb2*, and *Ncoa2* showed higher co-accessibility in primary infection than in reinfection (Figure 2D, bottom).

### Identification and characterization of B cell subpopulations in the murine spleen

Immune control of blood-stage parasitemia relies critically on B cells, the main effectors of humoral immunity^2^. As noted above, the splenic B cell compartment expanded sharply after the first parasite exposure and then contracted slightly after the second exposure (Figure S2F). However, changes in B cell frequency alone cannot explain the improved immunity observed upon reinfection, suggesting that qualitative remodeling of B cell phenotypes and functions also contributes to the establishment of immunological memory.

To dissect this compartment in more detail, we reanalyzed the 38,995 cells annotated as B cells based on *Cd19*, *Ms4a1*, and *Cd79* expression. Unsupervised clustering identified 13 transcriptionally and epigenetically distinct B cell subpopulations. UMAP projection revealed four major compartments: naïve/mature B cells, GCBs, memory B cells, and plasmablasts (Figure 3A). Subpopulations were annotated using canonical markers and DEGs specific to each cluster (Figure S3A, Table S1).

**Figure 3.**
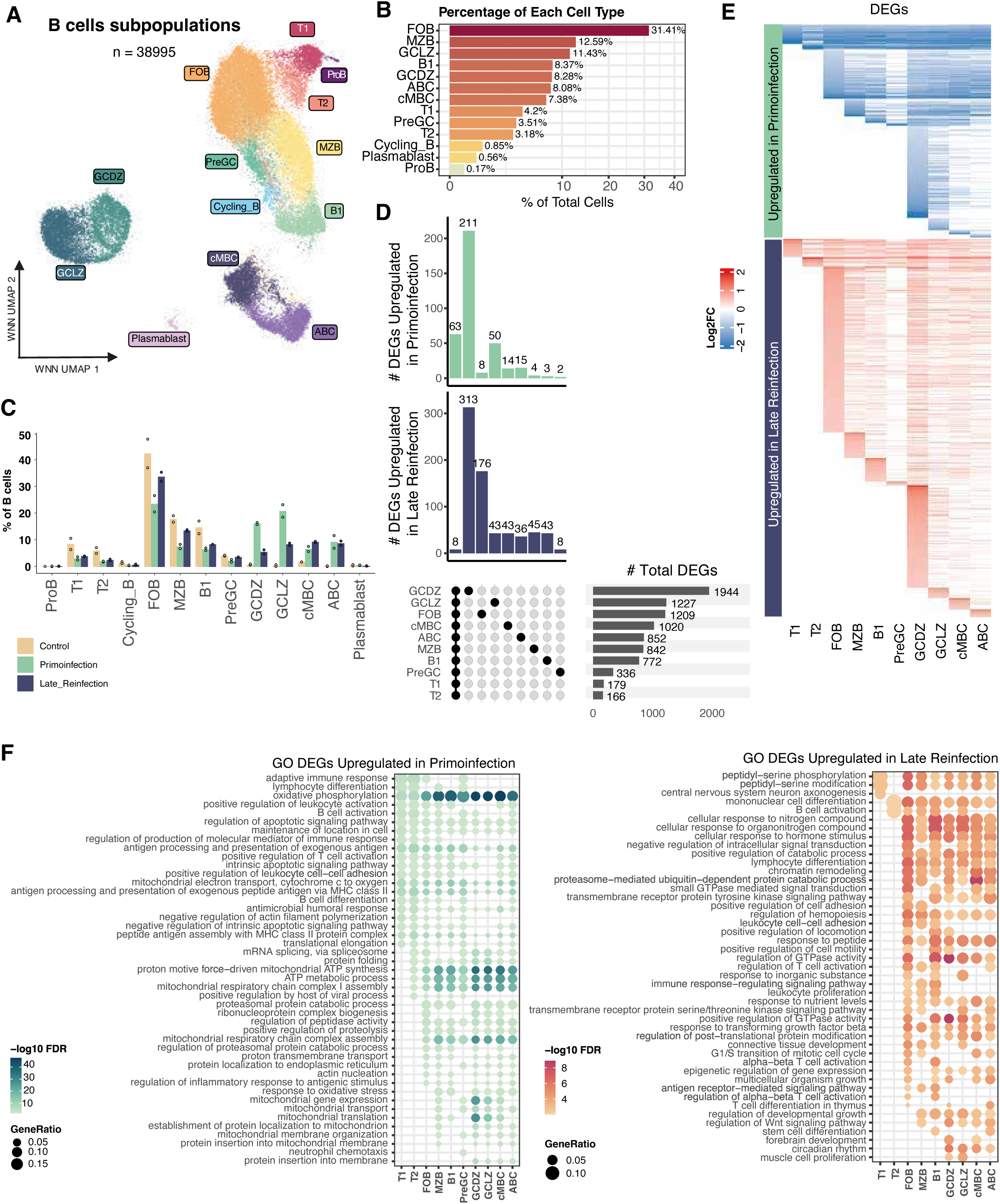
Transcriptional remodeling of B cell subpopulations upon Py17XNL infections. (A) WNN-integrated UMAP of the 38,995 B cells. Color indicates the B cell subpopulation to which each cell belongs. (B) Percentage of each B cell subpopulation relative to the total number of B cells in the atlas, ordered from most to least abundant. The color gradient indicates the percentage of cells. (C) Barplot showing the percentage of B cells in individual mice, grouped by experimental condition. Each bar represents the mean percentage for the two mice within the same infection group, while circles indicate individual values. (D) UpSet plot showing the overlap and exclusivity of DEGs when comparing late reinfection (LReinf) versus primary infection (Prim). (E) Heatmap showing log2FC values for DEGs when comparing reinfection versus primary infection in each subpopulation. (F) Functional enrichment analysis of GO Biological Process. Bubble plots showing the top 10 most enriched GO biological process terms in each B cell subpopulation among DEGs associated with primary infection (left) and reinfection (right). Circle size represents the proportion of DEGs annotated to each enriched term relative to the total number of DEGs, and the color gradient indicates −log10(FDR).

The largest compartment corresponded to naïve and mature B cells (64% of all B cells; Figure 3B), comprising eight subpopulations. Within this group, we identified a small progenitor B cell (ProB) population (n = 65), characterized by high *Ighm* and low *Ptprc* expression, and differential expression of *Sox4*, a TF implicated in early B cell (ProB) differentiation and survival^37^ (Table S1). Two transitional B cell subsets were also detected: T1 (n = 1,636), expressing *Ighm* and lacking *Fcer2a* (*CD23*), and T2 cells (n = 1,240), expressing *Ighm*, *Ighd*, and *Fcer2a* (Figure S3A). Among mature B cells, we resolved three classical subsets: follicular B cells (FOB), marginal zone B cells (MZB), and B1 cells. FOB cells (n = 12,249; 31% of all B cells; Figure 3B) were defined by *Ighd* and *Fcer2a* expression and low *Cr2* (*CD21*). MZB cells (n = 4,911) exhibited high *Cr2* together with *Pi3k4*, *Cd36*, and *Atxn1* (Table S1), consistent with their role in rapid responses to blood-borne antigens. B1 cells (n = 3,264) showed a distinctive profile with low *Ptprc*, high *Ighm*, low *Ighd*, and expression of *Il1ra5*, *Cacna1e*, and *Atxn1* (Table S1). In addition, clustering resolved recently described pre-germinal center (PreGC) B cells (n = 1,368; Figure S3B)^38,39^, as well as a cycling B cell subset (n = 331) expressing proliferation markers such as *Mki67* and *Top2a*.

The GCB compartment, defined by *Aicda* expression, accounted for ∼20% of B cells (Figure 3B, S3A) and contained two well-defined subpopulations: dark zone-like (GCDZ; centroblasts; n = 3,227), with a high proportion of proliferating cells (Figure S3C), and light zone-like (GCLZ; centrocytes; n = 4,458). These subpopulations showed the expected differential marker pattern^40^: GCDZ cells expressed high *Cxcr4* and low *Cd83* and *Cd86*, whereas GCLZ cells displayed low *Cxcr4* and high *Cd83* and *Cd86* (Figure S3D).

Memory B cells, defined by *Cd44, Ccr7* and *Cd80* expression, were subdivided into classical memory B cells (cMBCs; n = 2,877) and atypical memory B cells, also referred to as age-associated B cells (ABCs; n = 3,149). ABCs were characterized by *Itgax* (Cd11c) expression and by marker DEGs including *Zeb2*, a TF essential for ABC development^41^ (Table S1). Finally, plasmablasts were identified as a distinct compartment enriched for genes involved in terminal differentiation and antibody secretion.

To identify putative regulatory mechanisms underlying B cell subpopulation identity, we inferred TF activities. Notable TFs with subtype-restricted activity patterns included NFATC1 (NFAT2), largely confined to MZB and B1 cells; MEF2B, POU2F1 (OCT1), and POU2F2 (OCT2), enriched in GCBs; TBX21 (T-bet), specifically active in ABCs; and IRF4 and XBP1, selectively associated with plasmablasts (Figure S3E).

### Divergent cellular and molecular responses of B cell subpopulations to successive infections

After identifying B cell subpopulations, we next examined how their relative abundance changed with exposure to Py17XNL (Figure 3C). We observed a marked increase in GCB subsets (GCDZ and GCLZ) and memory B cells (cMBCs and ABCs) in infected samples, whereas in uninfected controls these populations represented only a minor fraction of the B cell compartment or were even absent, as in the case of ABCs. Conversely, naïve and mature B cell subsets, including T1, T2, cycling B cells, FOB, MZB, B1, and PreGC, were proportionally enriched in control samples, decreased sharply during primary infection, and partially recovered during reinfection, without returning to pre-infection levels. This pattern indicates a sustained imprint of infection on the composition of the splenic B cell pool.

When comparing infected groups directly, clear differences emerged as a function of the number of parasite exposures (Figure 3C). Both GCB subsets (GCDZ and GCLZ) were more abundant during primary infection and decreased by more than 50% in reinfection. In contrast, cMBCs increased in frequency during reinfection relative to primary infection, whereas ABCs remained at similar proportions in both conditions. Overall, these changes are consistent with a shift from a GC-dominated response during primary infection towards a memory-enriched B cell compartment upon reinfection, in line with more efficient recall responses after repeated parasite exposure.

We next asked whether repeated exposure to Py17XNL alters transcriptomic programs and chromatin accessibility within B cell subpopulations. Differential analyses were performed independently for each modality, focusing on pairwise comparisons between mice experiencing a single exposure (primary infection) and those with two exposures (reinfection). At the transcriptional level, we identified 2,761 DEGs overall between primary infection and reinfection across all B cell subpopulations (FDR < 0.05, |log2FC| > 0.5). The largest numbers of DEGs were detected, in descending order, in GCBs, FOB cells, and memory B cells (Figure 3D). In most subpopulations, DEGs were predominantly upregulated in reinfection, consistent with a more robust transcriptional activation associated with the secondary response (Figure 3E). Moreover, DEGs with higher expression in primary infection were broadly shared across multiple subpopulations, suggesting relatively similar transcriptional programs between subsets, whereas in reinfection overlap was reduced, suggesting more subpopulation-specific transcriptional responses (Figure 3E).

Several subpopulations showed substantial numbers of exposure-specific DEGs (Figure 3D). GCDZ cells displayed 524 unique DEGs, including histone variant genes such as *Hist1h1a*, *Hist1h3a*, and *Hist1h4a* overexpressed in primary infection, and apoptosis-related genes such as *Birc6* and *Trim17* overexpressed in reinfection. FOB cells exhibited 184 unique DEGs, including *Stat4*, *Stat6, Il21r*, and *Cxcr5*, all upregulated in reinfection. In addition, we identified a core set of 71 DEGs common to all B cell subpopulations (Figure 3D). Strikingly, only eight of these were overexpressed in reinfection, despite the strong global transcriptional activation observed in that condition, whereas the remaining 63 were more highly expressed in primoinfection. The expression patterns of these shared genes by subpopulation and condition are shown in Figure S3F. Genes enriched in primary infection included classical B cell genes involved in antigen presentation (*Cd74, B2m, H2-Aa, H2-Eb1, H2-Ab1*), BCR components (*Cd79a, Cd79b*), immunoglobulin genes (*Igkc, Iglc1, Iglc2, Iglc3*), mitochondrial function (*Cox4i1, Cox6c, Cox8a, Atp5e, Ndufa1*), and cytoskeleton/signaling regulators (*Tmsb4x, Tmsb10, Cfl1, Crip1*). Among the eight genes overexpressed in reinfection, we noted *Jarid2*, an epigenetic regulator that controls plasticity and cell fate through gene silencing^42–44^.

Over representation analysis (ORA) revealed distinct transcriptional programs associated with primary infection versus reinfection across B cell subpopulations (Figure 3F). DEGs upregulated in primary infection were enriched for mitochondrial metabolic processes, including cellular respiration, oxidative phosphorylation, and mitochondrial biogenesis, consistent with an early metabolic adjustment phase required for activation and expansion (Figure 3F, left). In contrast, reinfection was enriched for GO terms related to B cell activation and differentiation, antigen processing, immune receptor signaling, and adaptive lymphocyte responses, indicating a more specialized and functionally mature B cell activation state after secondary exposure (Figure 3F, right). Notably, enrichment for B cell activation-related genes was observed in both infection states (Figure 3F), suggesting that this pathway is engaged in primary and secondary responses but via differential gene usage.

Using a similar framework, we performed differential chromatin accessibility analysis to identify DARs between primary infection and reinfection. Although numerous accessible regions with nominal p-values < 0.05 were detected in B cell subpopulations (Figure S3G, top), only a small fraction passed an FDR < 0.2 (Figure S3G, bottom). This suggests that chromatin accessibility differences between primary and secondary infection in B cell subpopulations are relatively subtle and lack strong statistical support at the genome-wide level.

### Heterogeneity and origin of memory B cells in the spleen of infected mice

Memory B cells are essential for establishing effective and durable humoral immunity. Their generation requires prior antigen encounter, B cell activation and, in many cases, class-switch recombination and somatic hypermutation, which enable production of higher-affinity immunoglobulins during secondary responses. These processes drive transcriptional changes that reshape the functional heterogeneity of the memory B-cell compartment.

To characterize this diversity after Py17XNL infection, we re-clustered the full set of memory B cells (n = 6,026), comprising cMBCs and ABCs. This analysis resolved four clearly distinct clusters (Figure 4A), primarily separated by memory subset identity (cMBCs versus ABCs) and, within each, by immunoglobulin isotype: unswitched (IgM⁺) versus class-switched (IgM⁻) cells (Figure 4B). Subpopulation identity was validated using cMBC- and ABC-specific gene signatures^41^, which clearly separated cMBC and ABC populations; while cMBC-IgM⁺ cells showed an intermediate marker profile between both (Figure 4B).

**Figure 4.**
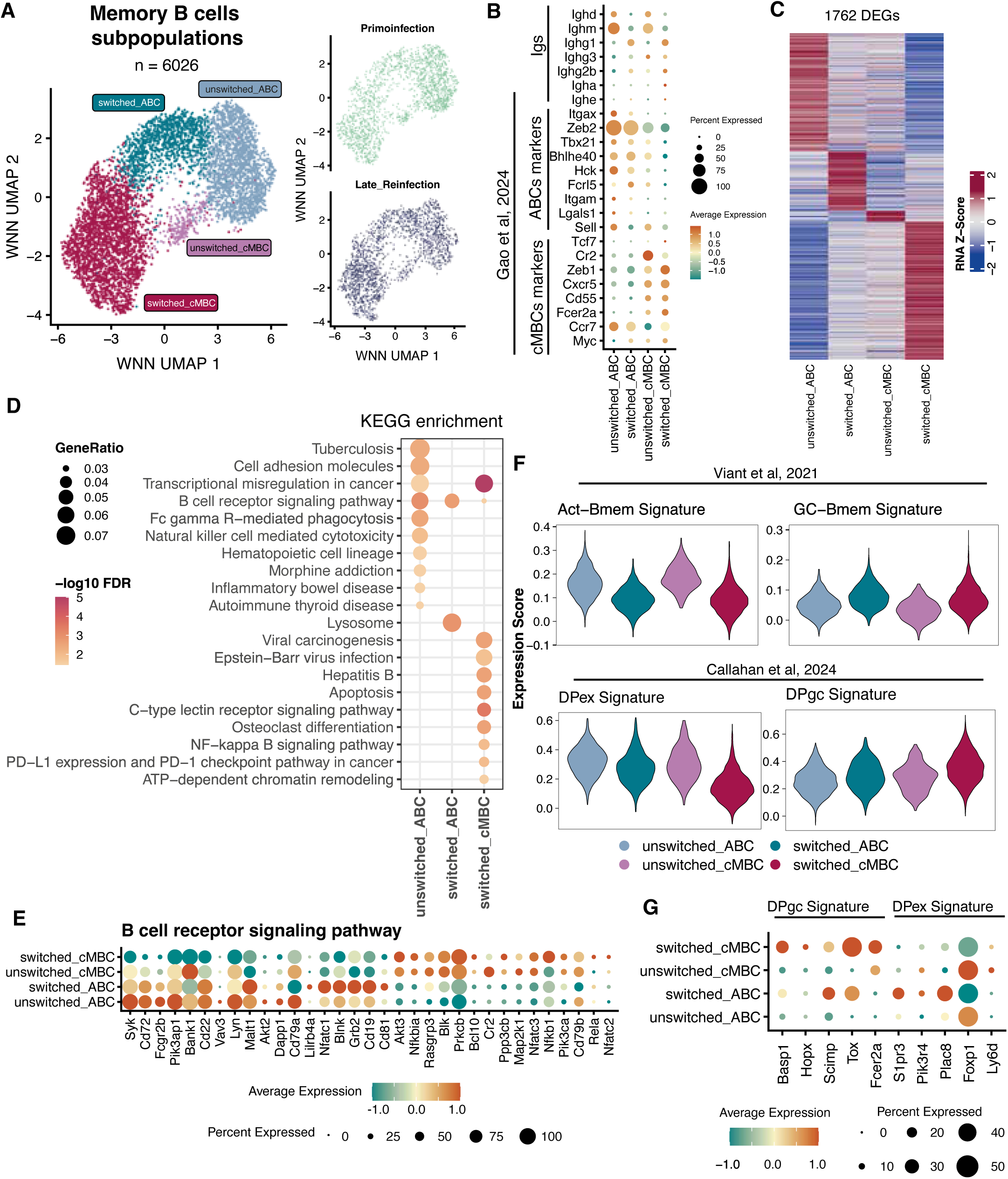
Characterization of memory B cell subpopulations. (A) UMAP projection of the identified memory B cell subpopulations, where each color represents a distinct group. On the right, UMAPs show memory B cells from primary infection and reinfection separately. (B) Expression of key marker genes in memory B cell subpopulations. (C) Heatmaps of differential gene expression showing the 1,762 DEGs across memory B cell subpopulations; each column represents a pseudobulk profile for a given subpopulation, and each row corresponds to a gene. The color scale indicates normalized expression values. (D) Top 10 enriched KEGG pathways for each subpopulation based on its marker genes; circle size is proportional to the ratio between DEGs and the total number of genes in each enriched term, and the color gradient indicates −log10(FDR). Circles highlight the subpopulation in which each term is enriched. (E) Mean expression of genes associated with BCR signaling in each subpopulation. (F) Violin plots showing expression scores for gene sets associated with extrafollicular versus GC origin for individual cells, grouped by subpopulation. (G) Mean expression of gene signatures from Callahan *et al.*^45^, stratified by GC-derived or extrafollicular origin. In panels B, E, and G, dot size indicates the proportion of cells expressing each gene, and color intensity represents standardized mean expression.

Differential expression analysis among the four subpopulations identified 1,762 DEGs (Figure 4C). BCR signaling was enriched in all subsets (Figure 4D), but with distinct transcriptional activation patterns: ABC-IgM⁺ cells showed higher expression of *Syk*, *Cd72*, *Fcgr2b*, *Pik3ap1*, *Bank1*, *Cd22*, *Vav3* and *Lyn*, whereas ABC-IgM⁻ cells expressed higher levels of *Nfatc1*, *Blnk*, *Grb2*, *Cd19* and *Cd81*. Both cMBC subsets (IgM⁺ and IgM⁻) expressed *Akt3*, *Rasgrp3*, *Blk*, *Prkcb*, *Bcl10*, *Cr2*, *Ppp3cb*, *Map2k1*, *Nfatc3*, *Pik3ca*, *Cd79b*, *Rela* and *Nfatc2*. The gene encoding the TF Nfkb1 was overexpressed exclusively in cMBC-IgM⁻ cells (Figure 4E). Consistent with this, cMBC-IgM⁻ cells were enriched for apoptosis, NF-κB signaling, PD-L1 expression and PD-1 checkpoint pathway in cancer, and ATP-dependent chromatin remodeling (Figure 4D).

To gain insight into whether each memory B cell subset preferentially reflects extrafollicular activation or GC reactions, we used mean expression of gene sets defined by Viant *et al.*^17^ and Callahan *et al.*^45^ (Figure 4F). In the first set of gene signatures^17^, we found that IgM⁺ cells showed higher expression of genes associated with extrafollicular-derived memory B cells (Act-Bmem), whereas IgM⁻ cells preferentially expressed GC-derived memory signatures (GC-Bmem). In the second set of gene signatures of CD80⁺PD-L2⁺ double-positive (DP) memory B cells^45^, cMBC-IgM⁻ cells were enriched for GC-derived DP signatures (DPgc), while the other three subpopulations were more enriched for extrafollicular DP signatures (DPex) (Figure 4F-G). These comparisons suggest differential weighting of GC-like versus extrafollicular-like programs across memory subsets rather than providing definitive lineage tracing.

We then assessed how memory B-cell heterogeneity changed between mice experiencing a single exposure (primary infection) versus two exposures (reinfection) to Py17XNL. The composition of memory B-cell subsets differed markedly between groups (Figure S4A): IgM⁺ cells predominated in primary infection samples, whereas IgM⁻ cells were more abundant in reinfection. Within memory B-cell subsets, ABC-IgM⁺ cells represented the largest fraction in primary infection, while cMBC-IgM⁻ cells dominated in reinfection.

At the transcriptomic level, cMBC-IgM⁻ cells showed the greatest number of changes between conditions, with 1,105 DEGs (FDR < 0.05, |log2FC| > 0.5) (Figure S4B), whereas cMBC-IgM⁺ cells displayed the fewest. ORA of DEGs with higher expression in reinfection (Figure S4C) revealed recurrent enrichment of epigenetic processes (chromatin remodeling), immune response pathways (regulation of lymphocyte activation and differentiation, antigen receptor-mediated signaling) and cell signaling (intracellular receptor-mediated signaling) across memory B-cell subsets. Both IgM⁺ groups were enriched for demethylation-related processes, and ABC-IgM⁻ cells additionally showed enrichment for immune response signaling and cytoplasmic pattern-recognition receptor pathways induced by viruses.

Using the memory B-cell marker gene sets detailed in Figure 4F, we performed GSEA for each of the four subpopulations with the same memory B-cell differentiation gene sets. Gene signatures associated with GC-derived memory B cells (GC-Bmem and DPgc) were significantly enriched among genes more highly expressed in primary infection samples (NES < 0, p-value < 0.05). Conversely, signatures corresponding to extrafollicular-origin memory B cells (Act-Bmem and DPex) showed positive enrichment (NES > 0) among genes more highly expressed in reinfection, although only the Act-Bmem set reached statistical significance (p-value < 0.05) in the cMBC-IgM⁻ subset (Figure S4D). These patterns support a shift towards extrafollicular-like transcriptional programs in reinfection, particularly within cMBC-IgM⁻ cells, but should be interpreted considering the limited number of biological replicates and the use of externally derived signatures.

Finally, we examined changes in TF activity between infection groups (reinfection versus primary infection) across memory B-cell subpopulations (Figure S4E). AP-1 complex activity (FOS, JUN, BATF, ATF3) was higher in all subpopulations from primary infection samples. No TF showed significantly enriched activity in reinfection, with the exceptions of the NF-Y complex (NFYC, NFYA, NFYB) in cMBC-IgM⁻ cells and ASCL2 in cMBC-IgM⁺ cells.

### Pseudo-temporal reconstruction of B cell differentiation lineages and regulatory dynamics during infection and reinfection

To explore splenic B cell differentiation dynamics during primary infection and reinfection with Py17XNL, we performed trajectory analysis using monocle3^46^ on the low-dimensional coordinates derived from scATAC-seq data (Figure 5A). Using chromatin accessibility patterns to order cells along pseudotime provides a more stable and predictive representation of cell states than gene expression alone, which can fluctuate more rapidly. Before trajectory inference, we excluded cycling B cells, due to their population heterogeneity, and plasmablasts, because of their low abundance and highly specialized transcriptomic and chromatin profiles.

**Figure 5.**
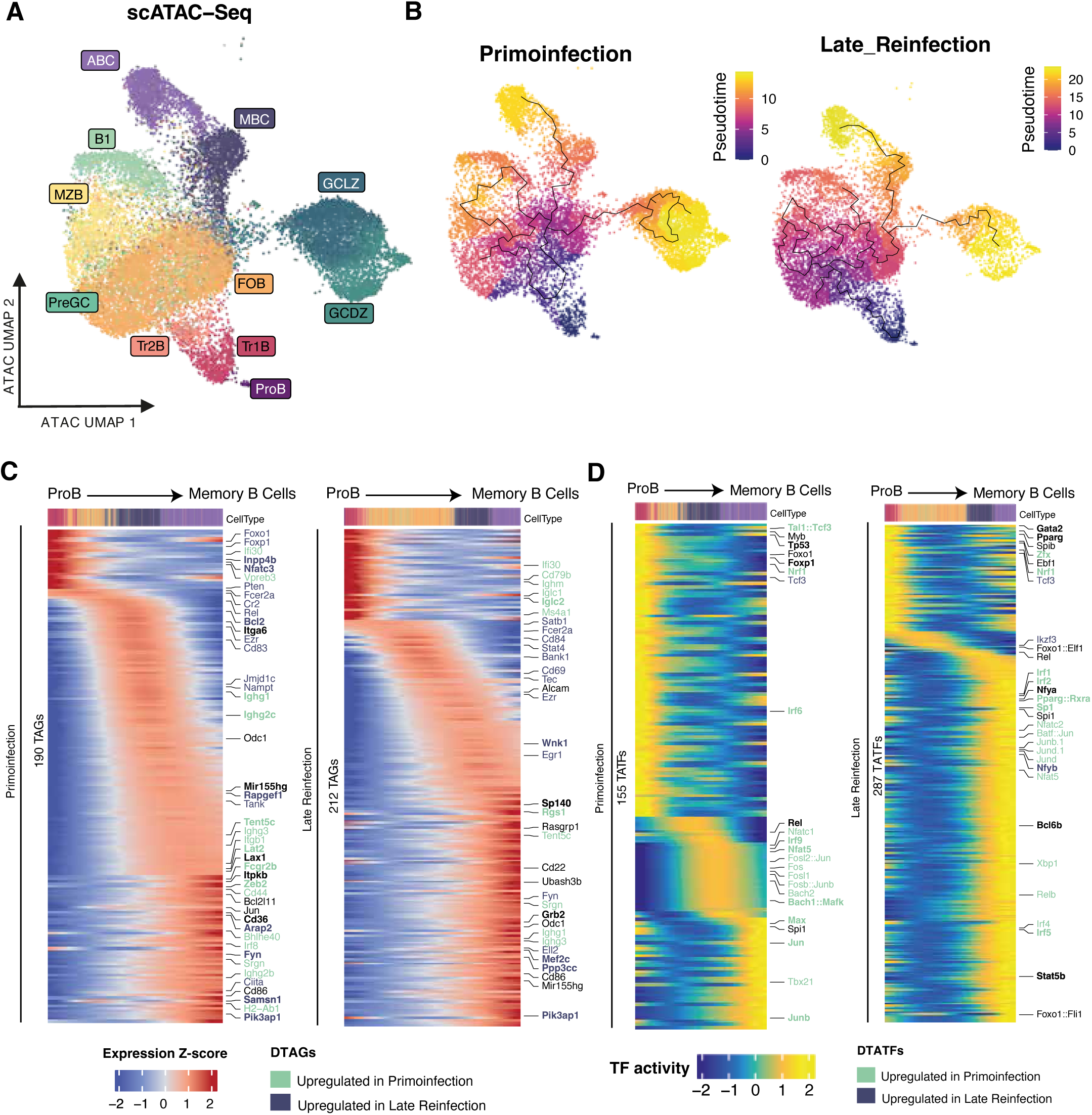
Dynamics of gene expression and TF activity along pseudotime of memory B cell trajectory. (A) UMAP of the scATAC-seq integration for the 38,995 B cells. Color indicates the B cell subpopulation to which each cell belongs. (B) UMAPs showing the results of trajectory analysis for B cells in primary infection (left) and reinfection (right). The color gradient represents the pseudotime value assigned to each cell, and the black line indicates the inferred trajectory. (C) Gene expression changes associated with memory B cell trajectories. Heatmaps showing genes whose expression changes along B cell trajectories inferred in primary infection (left) and reinfection (right). Genes in bold are trajectory-specific genes when comparing trajectories within the same condition. Genes in green represent those with higher expression in primary infection samples, and genes in blue denote those with higher expression in reinfection samples. The heatmap color gradient indicates standardized expression values along the trajectory. Cells are ordered by pseudotime. TAGs (trajectory-associated genes) and DTAGs (TAGs whose expression changes significantly in both contexts, primary infection and reinfection) are indicated. (D) TF activity changes associated with memory B cell trajectories. Heatmaps showing TFs whose activity changes along B cell trajectories in primary infection (left) and reinfection (right). TFs in bold are trajectory-specific when comparing trajectories within the same condition. TFs in green have higher activity in primary infection, whereas TFs in blue have higher activity in reinfection. The heatmap color gradient indicates TF activity values along the trajectory. Cells are ordered by pseudotime.

We defined ProB cells as the root population, which enabled us to clearly resolve three differentiation trajectories under each infection condition (Figure 5B). These trajectories converged toward GCBs, memory B cells, or mature B cells (FOB, MZB, and B1). Pseudotime values reflected progression along each lineage and differed between infection conditions: reinfection trajectories spanned a broader pseudotime range (0-24), indicating a more extended differentiation continuum, whereas primary infection trajectories covered a shorter range (0-15), consistent with a faster or more constrained differentiation process (Figure 5B).

Once trajectories were defined, we next examined changes in gene expression and TF activity along each lineage of memory B cells (Figure 5C, D). We focused on genes and TFs whose activity varied significantly with pseudotime, thereby inferring their potential contribution to shaping lineage progression without explicitly contrasting conditions. Overall, the number of trajectory-associated genes (TAGs) was higher in reinfection than in primary infection across B cell lineages (FDR < 0.05, mean log2FC > 0.5), except for GCB trajectories (Figure 5C).

TAGs formed expression modules with distinct pseudotemporal patterns (Figure 5C) across all trajectories. Early pseudotime was characterized by high expression of broadly B cell-associated genes such as *Foxo1* and *Ifi30*. At intermediate pseudotime, genes including *Satb1*, *Fcer2a*, and *Cr2* were more prominently expressed, suggesting roles in activation and modulation of immune responses during intermediate states. At late pseudotime, genes such as *Cd44*, *Cd86*, and *Ell2* showed increased expression, consistent with roles in terminal differentiation and mature B cell function. Lineage-specific TAGs were also identified.

We then focused on TAGs whose behavior differed between primary infection and reinfection along these trajectories (DTAGs; Figure 5C). In the memory B cell lineage, *Zeb2*, *Rgs1*, and *Srgn* were more highly expressed in primary infection, whereas *Nfatc3*, *Bcl2*, *Fyn*, and *Wnk1* showed higher mean expression in reinfection.

We also assessed changes in TF activity along pseudotime and between infection conditions (Figure 5D). A set of TFs associated with intermediate and late pseudotime stages, including members of the AP-1 complex (FOS, FOSL2, JUN, JUNB), BACH2, and interferon-regulated TFs (IRF9, IRF4, IRF8), displayed higher activity in primary infection. Lineage-specific TFs were also evident: BCL6 and SPI family members (SPI1, SPIB, SPIC) were enriched along the GCB trajectory, with SPI family TFs showing higher activity in reinfection.

### Inference of gene regulatory networks in B cells during primary infection and reinfection

To investigate the gene regulatory programs that define B cell states in response to *Plasmodium* infection, we inferred gene regulatory networks (GRNs) by integrating single-cell chromatin accessibility and gene expression data. We used an adapted version of GRaNIE^47^, computing GRNs independently for each previously identified trajectory branch, thereby aiming to capture regulatory interactions specific to B cell lineages and infection conditions.

The resulting GRNs contained between 68 and 94 TFs, each correlated with 325-5,371 accessible peaks and 401-2,800 unique target genes, with a median of 1,087 regulatory interactions per GRN (Figure 6A). The most connected network (13,720 interactions) corresponded to the GCB trajectory in primary infection. Between 20-25% of TFs were unique to a given GRN, with the remainder shared across multiple networks (Figure 6B). Four TFs (BACH2, FOSL2, JUNB, and SPIB) were present in all GRNs, whereas others were condition specific: FOSL1, NFKB, and RBPJ were unique to primary infection GRNs, while EGR1, ELF4, ETS1, and PRDM9 were exclusive to reinfection GRNs.

**Figure 6.**
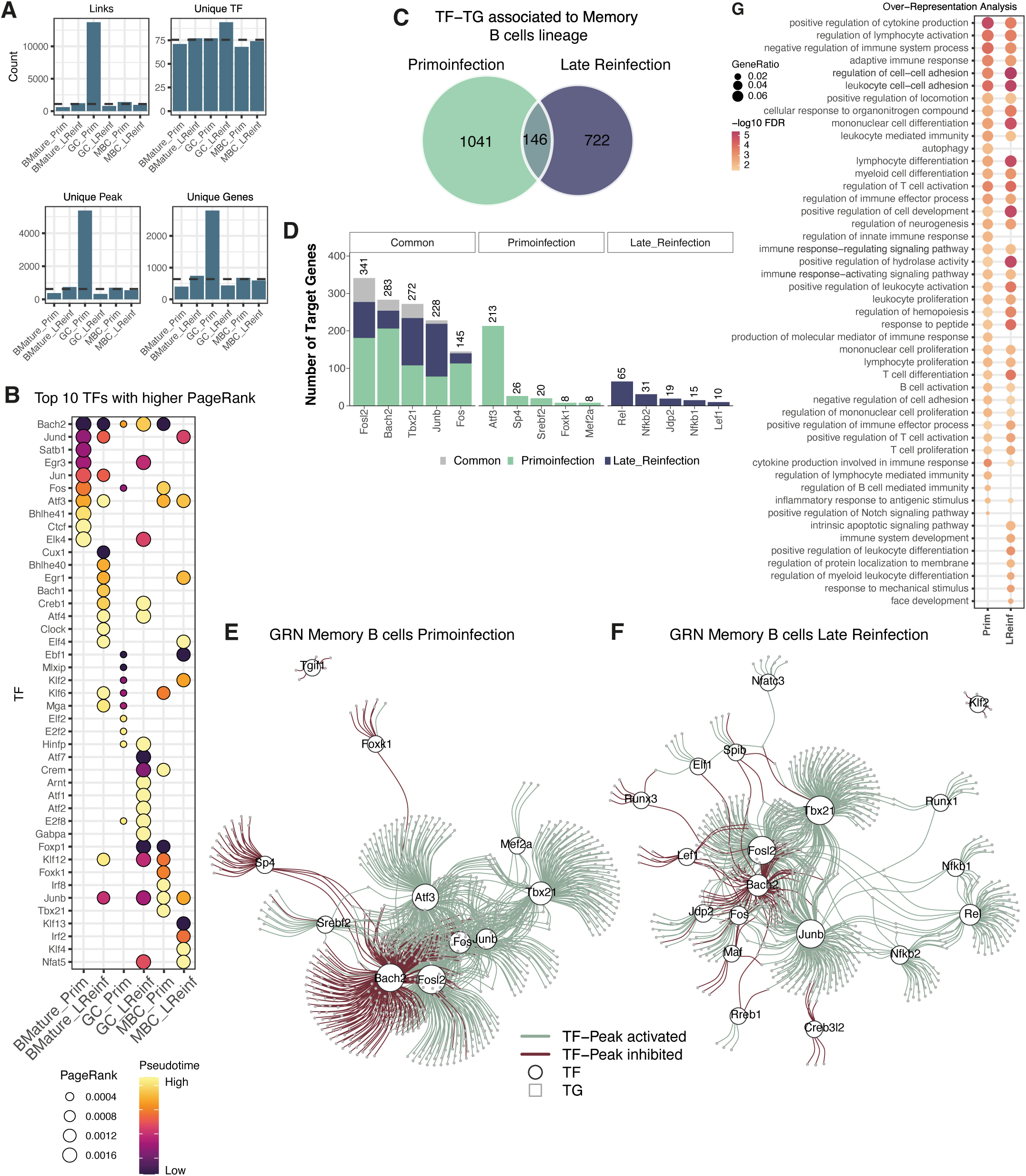
GRNs associated with splenic B cell lineages. (A) Number of TF-peak-target interactions, TFs, peaks, and unique target genes identified in each network. The dashed line indicates the median of each metric. (B) Bubble plot showing the top 10 TFs ranked by PageRank in each GRN. The color gradient represents the pseudotime at which each TF reaches its maximum expression, and bubble size reflects the PageRank value, a nodal centrality score. MBC denotes memory B cells, Prim indicates primary infection, and LReinf indicates reinfection. (C) Regulatory networks associated with memory B cells. Venn diagram showing overlap in TF-target interactions between GRNs inferred from primary infection and reinfection. (D) Number of target genes per TF, showing the top 5 TFs with the highest number of targets, both shared and condition specific. (E/F) Representative GRNs for primoinfection (E) and reinfection (F). White circles represent TFs and small grey circles represent target genes. TF name and node size reflect degree centrality in each GRN, with larger circles and labels indicating higher centrality. Green edges represent positive correlations (R>0), and red edges represent negative correlations (R<0). (G) Top 30 enriched GO biological process terms based on all TFs and target genes in each GRN. Circle size is proportional to the ratio between TFs/targets and the total number of genes in each enriched term, and the color gradient indicates −log10(FDR). Circles highlight the GRN in which each term is enriched.

To prioritize TFs with the greatest topological importance, we computed PageRank centrality for TFs whose expression or activity varied along the trajectory corresponding to each GRN and selected the top 10 TFs per network (Figure 6B). These highly connected TFs represent nodes with strong inferred regulatory influence over target genes in specific B cell states. Although many TFs appeared across multiple GRNs, both their degree centrality and the pseudotime at which they reached maximal expression varied markedly. For example, BACH2, present in nearly all GRNs, showed relatively low connectivity in GCB lineage networks but substantially higher connectivity in the memory B cell GRN from primary infection. Condition-specific differences were also evident: JUNB, shared by both memory B cell GRNs, exhibited higher centrality in reinfection, suggesting dynamic and condition-dependent transcriptional reprogramming (Figure 6B).

To further dissect condition-specific regulation, we focused on the GRNs derived from the memory B cell lineage in primary infection and reinfection. These two networks shared only 146 TF-target interactions (Figure 6C), corresponding to 14% of the primary infection network and 20% of the reinfection network. This low overlap suggests extensive rewiring of regulatory circuits controlling memory B cell differentiation and maintenance as a function of the number of parasite exposures, consistent with more specialized transcriptional and epigenetic adaptation during secondary responses.

Both networks contained TFs with known roles in memory B cell identity, including TBX21, BACH2, and FOSL2, but their target gene sets differed markedly between conditions (Figure 6D). Additional TFs were unique to a single condition: ATF3 and SP4 appeared only in the primary infection GRN, whereas REL, NFKB1, and NFKB2 were detected exclusively in reinfection. This divergence in active TFs suggests that memory B cells do not simply maintain a fixed core of regulators but instead recruit additional condition-specific TFs after repeated pathogen exposure from our inferred networks. These differences were also reflected in GRN topology (Figure 6E,F): the primary infection GRN was dominated by BACH2, FOSL2, ATF3, and TBX21, whereas the reinfection GRN was dominated by JUNB, FOSL2, and TBX21 together with REL and NFKB2.

Functional ORA of each GRN, using the full set of TFs and targets, revealed broadly similar GO enrichments across conditions, including cytokine regulation, adaptive immune responses, lymphocyte differentiation, and B cell activation (Figure 6G). In primary infection, BACH2, TBX21, ATF3, FOS, and FOSL2 regulated many B cell activation-related genes, consistent with their higher activity in this group (Figure 5D). In contrast, some biological processes were uniquely enriched in one condition. The primary infection GRN showed enrichment for autophagy, regulation of innate immune responses, and Notch signaling, whereas the reinfection GRN was enriched for intrinsic apoptotic signaling, leukocyte differentiation, and responses to mechanical stimuli (Figure 6G).

To identify TFs regulating genes associated with distinct memory B cell differentiation routes, we constructed subnetworks centered on gene sets previously linked to extrafollicular- versus GC-derived memory^17,45^ (Figure S5). In both conditions, more target genes were associated with GC-derived programs (GCBmem and DPgc) than with extrafollicular programs, particularly those defined by Act-Bmem. Among TFs regulating the largest number of these targets TBX21, FOSL2, BACH2, and ATF3 in primary infection, and JUNB, FOSL2, BACH2, TBX21, and REL in reinfection. Some TFs showed preferential association with specific memory differentiation paths: ATF3 was connected to GCBmem genes; BACH2, NFKB1, and JUNB were characteristic of DPgc; and REL was linked to extrafollicular (Act-Bmem) differentiation.

## Discussion

*Plasmodium* is the protozoan parasite that causes malaria, an endemic disease in tropical and subtropical regions that remains a major cause of mortality, particularly in children under five years of age (WHO, 2023). Unlike most infectious diseases, protective immunity to malaria in endemic populations develops only after repeated parasite exposure and is typically non-sterilizing, failing to fully prevent subsequent infections^2,3^. This atypical pattern of immunity raises fundamental questions about how repeated blood-stage infections progressively reshape lymphocyte differentiation programs and fine-tune effector functions^3,5^.

To model this scenario experimentally, we selected Py17XNL because it causes a self-resolving infection with relatively low parasitemia (<30%), allowing spontaneous parasite clearance and the development of protective immunity to reinfection^48–51^, in contrast to *P. chabaudi chabaudi*, which can reach parasitemia of ∼50% and establish chronic infections^52–54^. In our hands, primary infection with Py17XNL led to peak of parasitemia around 13 dpi and complete parasite clearance from peripheral blood by ∼18-21 dpi. whereas reinfected mice did not develop microscopically detectable parasitemia, consistent with the induction of robust protection and with previous reports using this model^55^. Notably, immunity induced by non-lethal Py17XNL can also protect against lethal strains^56^, underscoring the relevance of this model for studying acquired anti-malarial immunity.

The spleen is a central site for blood-stage parasite clearance and for initiation and maintenance of anti-malarial humoral immunity^6–9^. Splenectomized mice infected with non-lethal *Plasmodium* strains fail to control parasitemia and show high mortality^57^, and splenectomized humans experiencing a first *P. falciparum* infection exhibit higher parasitemia, impaired clearance, and increased risk of severe disease and death^58,59^. *Plasmodium* infection induces splenomegaly and profound remodeling of splenic architecture, including *de novo* GC formation, distortion or loss of marginal zone and T cell compartments, and marked changes in cellular composition^6,7,9,11^. The extent and nature of this disruption depend on both host and parasite strain: lethal infections often show disorganized or ineffective GCs^1,60^, whereas non-lethal *P. chabaudi chabaudi* maintains GC formation despite loss of marginal zone B cells and T cell zone disorganization^61^. Murine models based on non-lethal strains such as Py17XNL recapitulate key features of human malaria, including self-resolving primary parasitemia and strong protection against subsequent challenge^12,13^, providing a tractable system to dissect how primary and recall responses are encoded within splenic lymphocyte compartments^14,15^.

Here we applied a multimodal single-cell approach to interrogate the molecular programs underlying immune memory to Py17XNL. By integrating scRNA-seq and scATAC-seq data, we characterized infection- and reinfection-associated changes in gene expression, linked them to chromatin remodeling, and inferred the activity of specific TFs across distinct immune cell types. This strategy aligns with recent advances in systems immunology, shifting from static enumeration of cell subsets toward a dynamic, regulatory view of how cellular states are encoded and reshaped during primary and recall responses. Our multiomic atlas of >50,000 splenic mononuclear cells from control, primary infection, and early and late reinfection expands previous single-cell studies focused on acute malaria^62,63^ or on reinfection after artemisinin treatment restricted to selected T or B cell subsets^14,64^, and to our knowledge provides the first integrated view of naturally resolved primary infection and reinfection in this model.

A first key observation from this atlas is that B cells dominate splenic remodeling during Py17XNL infection. Among the 13 major splenic cell types identified, B cells were the most abundant, accounting for nearly 70% of mononuclear cells, substantially higher than the 33-50% reported in steady-state conditions^65,66^. This over-representation is consistent with previous murine studies that report robust splenic B cell expansion across *Plasmodium* strains^67–69^. However, changes across B cell subsets were not uniform. Naïve and mature follicular and marginal zone B cells decreased after primary infection and only partially recovered after reinfection, whereas GC B cells (dark zone centroblasts and light zone centrocytes) markedly expanded during primary infection and remained elevated during reinfection. This pattern agrees with previous reports showing GCs initiating around 7 dpi, peaking around 20 dpi, and persisting up to ∼60 dpi^60^, and suggests that early GC formation after first exposure is critical to establish a response able to limit parasite growth and support protection against subsequent infections.

Within the memory compartment, we detected expansion of both classical memory B cells (cMBCs) and atypical/age-associated B cells (ABCs) after primary infection, indicating that a single Py17XNL exposure is sufficient to generate diverse memory subsets, as previously shown^70^. cMBCs mediate accelerated and more effective responses upon repeated pathogen exposure^16,71^. ABCs have been previously described in malaria as well as in other chronic infections or in autoimmune diseases^3,72–74^. Although initially considered dysfunctional, ABCs are now recognized as GC-derived, functionally active, and capable of responding to re-exposure, contributing to control of chronic infections despite unclear longevity^75–77^. In our model upon reinfection, cMBCs continued to increase, while ABC numbers remained relatively stable, suggesting that cMBCs have greater proliferative or expansion capacity during secondary responses and likely contribute more actively to protection, whereas ABCs may perform more specialized, numerically stable functions. After primary infection, both isotype-switched (IgM⁻) and unswitched (IgM⁺) memory B cells were already present, consistent with rapid generation and long-term persistence of both subsets^25^. Reinfection reshaped this balance: IgM⁺ memory B cells predominated in primary infection, whereas IgM⁻ memory B cells were more abundant during reinfection. A relative decline of IgM⁺ memory cells after repeated exposure has been reported^25^, suggesting contraction of this subset during late or secondary phases. Nevertheless, IgM⁺ memory B cells remain functionally critical, as they generate most antibody-secreting cells upon reinfection and carry somatic mutations consistent with high affinity^25^. Our data therefore support a model in which numerically reduced IgM⁺ memory B cells contribute disproportionately to early secondary humoral responses, while IgM⁻ cMBCs accumulate and may support sustained, class-switched immunity. Moreover, it should be noted that the transcriptional profiles of cMBCs and ABCs in our atlas closely matched previously reported phenotypes, as the *Zeb2*^77^, supporting accuracy of our annotations. *Zeb2*, predominantly expressed in ABCs, has been linked to their development together with TBX21^77^. Consistently, chromatin accessibility analysis using chromVAR revealed specific enrichment of TBX21 motifs in ABCs. In contrast, *Zeb1*, more highly expressed in cMBCs, has been implicated in GC formation and in the establishment of memory B cells^78,79^.

A limitation of our study is the underrepresentation of plasma cells. We detected only ∼200 plasmablasts across all samples, precluding detailed analysis. This number is lower than in other splenic single-cell studies of malaria infection^62,64^, likely because our sampling time points occurred after the peak of the acute response. Previous work shows plasmablasts peaking around 10 dpi, decreasing by ∼20 dpi, and nearly disappearing between 28-30 dpi^25,61,80^. We also did not observe a marked plasmablast expansion after reinfection, consistent with some reports^81^ but contrasting with others documenting strong secondary plasmablast bursts^25,26^. The limited plasmablast expansion in our model may reflect rapid, efficient recall responses driven by pre-existing memory, reducing the need for large de novo plasmablast bursts. However, we cannot determine whether these plasmablast responses are short- or long-lived, as this would require parallel analysis of blood and bone marrow.

At the molecular level, repeated parasite exposure reshaped the transcriptional programs of splenic B cells. Differential expression analysis identified >2,700 DEGs between primary infection and reinfection, indicating substantial plasticity across B cell subpopulations. Most DEGs were upregulated in reinfection, consistent with a more robust and specialized transcriptional activation during secondary responses, particularly in GCBs, FOB cells, and memory B cells, the subpopulations with the largest DEG counts. In primary infection, upregulated genes were enriched in pathways associated with early activation and expansion, including antigen presentation, adaptive responses, and energetic metabolism. Among the 63 DEGs shared by all B cell subpopulations, we identified BCR components (*Cd79a*, *Cd79b*), immunoglobulin genes, mitochondrial genes, and cytoskeleton-related factors, consistent with a proliferative, activation-focused program. BCR signaling and cytoskeletal remodeling are essential for effective B cell activation^82,83^, and enrichment of oxidative phosphorylation, mitochondrial, and ribosomal genes suggests metabolic reprogramming to meet the energetic demands of proliferation, in line with previous studies^84,85^.

In contrast, reinfection was dominated by DEGs associated with intracellular signaling, lymphocyte activation and differentiation, and epigenetic regulation, including chromatin remodeling. Notably, *Jarid2* was consistently overexpressed across all B cell subpopulations. *Jarid2* functions as a transcriptional repressor that recruits the Polycomb repressive complex 2 (PRC2) to target genes^86^ and has been linked to inhibition of cell proliferation^87,88^. This matches our observation of reduced proliferative activity during reinfection compared with the more proliferative primary response, suggesting that *Jarid2* may help limit clonal expansion at later stages through epigenetic repression. PRC2 is a core Polycomb group complex that mediates stable gene repression via post-translational histone modification, most notably H3K27 trimethylation (H3K27me3), which promotes chromatin compaction and durable transcriptional silencing^89–93^. In B cells, coordinated activity of PRC1 and PRC2 regulates transitions between resting and proliferative states in GCs and controls differentiation, proliferation, and maintenance of B cell identity^94,95^. Together, these findings suggest that repeated *Plasmodium* exposure engages epigenetic modules that restrain proliferation while preserving a poised, memory-competent state.

GSEA showed that GC differentiation-associated gene signatures predominated in primary infection across all memory subsets, whereas reinfection, especially in IgM⁻ cMBCs, was enriched for extrafollicular-like programs. Together with previous work indicating that IgM⁺ memory B cells preferentially re-enter GCs while IgM⁻ memory B cells more often differentiate directly into plasma cells via extrafollicular routes^96–99^, and with recent data implicating extrafollicular-like memory B cells in improved protection upon reinfection in malaria models^15^, our results suggest that repeated Py17XNL exposure biases the memory compartment toward IgM⁻ extrafollicular-like cells poised to mount rapid secondary antibody responses without extensive GC re-entry.

GSEA revealed enrichment of GC differentiation-associated genes predominantly in primary infection across all memory subsets. In contrast, reinfection showed increased expression of genes linked to extrafollicular differentiation, most prominently in IgM⁻ cMBCs. Together with previous work indicating that IgM⁺ memory B cells preferentially re-enter GCs while IgM⁻ memory B cells more often differentiate directly into plasma cells via extrafollicular routes^96–98,100^, and with recent data implicating extrafollicular-like memory B cells in improved protection upon reinfection in malaria models^15^, our results suggest that repeated Py17XNL exposure biases the memory compartment toward IgM⁻ extrafollicular-like cells poised to mount rapid secondary antibody responses without extensive GC re-entry.

B cell diversity emerges from a hierarchical hematopoietic program in which bone marrow progenitors generate immature B cells that mature in secondary lymphoid organs into naïve, FOB or MZB cells and, upon activation, differentiate, with or without passing through germinal centers, into memory B cells or plasma cells^101^. This hierarchy is overlaid by marked functional plasticity, driven by extrinsic cues, lineage-defining TFs and large-scale transcriptional reprogramming, allowing transitions from immature precursors to specialized effector and memory states^102–106^. In this context, our trajectory inference on scATAC-seq data^107,108^ ordered splenic B cells from Py17XNL-infected mice along three main differentiation paths converging on GCBs, memory B cells or mature B cells

Along these trajectories, combined analysis of gene expression and TF activity identified lineage- and context-specific regulators. We observed marked differences between primary infection and reinfection in TF activity patterns. Members of the AP-1 complex (FOS, FOSL1/Fra-1, FOSL2/Fra-2, JUN, JUNB) displayed higher activity in primary infection across all three trajectories and were preferentially engaged at intermediate and late pseudotime. Given their established roles in regulating proliferation, differentiation, and apoptosis^109^, and in B cell maturation^110^, elevated AP-1 activity during primary infection likely supports clonal expansion, transcriptional remodeling, and effector differentiation needed to establish memory. In contrast, its reduced activity in reinfection is consistent with a more stable, specialized transcriptional state dominated by pre-existing memory B cells and biased toward rapid antibody production rather than de novo proliferative expansion.

Joint transcriptomic and epigenomic analysis revealed temporal coupling between TF expression and accessibility at their binding motifs. For some TFs, such as ATF3, changes in expression preceded increased accessibility at their cognate sites, suggesting a pioneer-like role in priming chromatin before target gene induction and biasing cells toward specific lineages by preconfiguring the epigenetic landscape^111^. In other cases, including TBX21, TF expression increased only after chromatin at its motifs was already accessible, consistent with a role in activating pre-open CREs rather than establishing accessibility per se. Together, these patterns illustrate how distinct TF modes of action contribute to the epigenetic remodeling and functional plasticity that differentiate primary from secondary B cell responses.

Inferring GRNs from single-cell data provides a powerful framework to dissect molecular mechanisms underlying cell-state transitions. Recent multiomic assays that jointly profile chromatin accessibility and gene expression have enabled GRN methods that explicitly integrate these layers^112–114^. Most approaches, however, infer a single global GRN from all cells in a dataset, maximizing TF-target coverage but potentially masking interactions restricted to specific states or conditions, such as those induced by infection.

Here, we adopted a more granular strategy, inferring GRNs independently for each infection condition within the monocle3-defined B cell trajectories using GRaNIE^47^, which integrates epigenomic (chromatin accessibility) and transcriptomic data to derive TF-peak-gene links. Applying GRaNIE per trajectory branch provided several advantages: (i) reduced biological heterogeneity by avoiding the pooling of transcriptionally divergent cell types, (ii) improved contextual specificity, allowing detection of regulators that are active only in particular developmental or immune states, and (iii) more interpretable networks, directly linked to defined lineage branches and infection stages. GRaNIE has been validated for cell type-specific GRNs in bulk^115,116^ and single-cell settings^117^, but typically after aggregating conditions and cell types to increase power, at the cost of specificity. In contrast, our trajectory-focused approach yields higher-resolution, condition-specific regulatory models by restricting inference to cells within a single branch.

Within this framework, GRNs inferred along the memory B cell trajectory highlighted TFs that modulate memory identity and recall responses after repeated parasite exposure with TBX21, FOSL2, and BACH2 emerging as central regulators. BACH2, a repressive TF that promotes memory B cell development while limiting terminal effector differentiation, has been linked to plasma cell differentiation when downregulated in class-switched (IgM⁻) memory B cells^118^. In line with this, we observed higher BACH2 expression, activity and regulon size in primary infection than in reinfection, suggesting that BACH2 helps maintain a less differentiated memory-favoring state during the primary response, whereas its downregulation in reinfection may facilitate rapid memory B cell-to-plasma cell conversion and an optimized secondary antibody response. Our GRNs further indicate that BACH2 represses a broad set of GC B cell differentiation genes, supporting a role in promoting extrafollicular-like memory programs during primary infection. In addition, the multiomic GRNs revealed a layered regulatory architecture in which ATF3 strongly controlled memory lineage genes in primary infection; ATF3 is a stress-inducible TF that modulates metabolism, immune responses and regulatory B cell function^119–121^ and can interact with the AP-1 complex^122^, suggesting potential synergy with AP-1 TFs preferentially activated in primary infection to enhance memory B cell activation and differentiation.

In contrast, REL (c-Rel) and NFKB2 (p100/p52), both members of the NF-κB family^123^ with well-established roles in B cells and GCBs^124–129^, emerged in our reinfection GRN, as hallmark regulators that control an expanded set of memory B cell target genes, consistent with a more complex and fine-tuned transcriptional program required for rapid, efficient secondary responses. These findings support a context-specific role for NF-κB family members in memory B cell reprogramming during reinfection, although targeted functional studies will be needed to define their precise contribution to recall fate and function.

## Methods

### Mouse model of sequential *P. yoelii* 17XNL infection

All animal procedures were conducted in accordance with EU Directive 2010/63, Spanish legislation (R.D. 53/2013), and were approved by the Animal Experimentation Ethics Committee of the Complutense University of Madrid (UCM). Experiments were performed in the animal facility of the Faculty of Veterinary Medicine at UCM. Female BALB/c mice (8-10 weeks old) were maintained under specific-pathogen-free conditions with a 12 h light/dark cycle, controlled temperature (22-24°C) and ∼50% relative humidity, with food and water provided ad libitum.

Infection was induced with the non-lethal Py17XNL strain, which causes self-limiting infections and allows the development of immunological memory. Mice were inoculated intraperitoneally with 1 × 10⁶ infected red blood cells (iRBCs) obtained from a BALB/c donor mouse infected with the same strain. Parasitemia and body weight were monitored every 2-3 days from the beginning of the experiment using tail vein blood smears stained with Wright eosin-methylene blue and quantified with Plasmoscore 1.3 software (Burnet Institute, Melbourne, Australia).

To assess memory responses, a subset of animals was reinfected with the same dose of iRBCs on day 34 post-infection (dpi), only when parasitemia was undetectable (0%) by microscopy. Mice were allocated to four experimental groups: (i) control (Ctrl), non-infected animals; (ii) primary infection (Prim), mice infected once and sacrificed at 34 dpi; (iii) early reinfection (EReinf), mice reinfected and sacrificed at 5 days post-reinfection (dpr; 39 dpi); and (iv) late reinfection (LReinf), mice reinfected and sacrificed at 15 dpr (49 dpi). Spleens were collected at each time point for single-cell analyses.

### Isolation of splenic mononuclear cells

Spleens were manually dissociated into ∼10 mm³ fragments in RPMI medium supplemented with 10% fetal bovine serum (FBS) and 5 µL DNase I and incubated for 5 min at room temperature. Tissue fragments were then gently triturated and passed through a 70 µm nylon strainer using a syringe plunger, washing the filter with 5 mL of cold RPMI + 10% FBS.

Cells were centrifuged at 500 g for 5 min at 4 °C, the supernatant was discarded, and the pellet was resuspended in 5 mL PBS at room temperature. Splenic mononuclear cells (lymphocytes and monocytes) were isolated by density gradient centrifugation using Ficoll-Paque (400 g, 20 min, room temperature). The mononuclear layer was collected and washed twice with cold RPMI + 10% FBS, centrifuging and discarding the supernatant each time.

The final cell suspension was resuspended in RPMI + 40% FBS. Cell numbers were determined in a Neubauer chamber after a 1:40 dilution in PBS and trypan blue staining to estimate viability. For cryopreservation, cells were resuspended in freezing medium (30% dimethyl sulfoxide [DMSO] in RPMI + 40% FBS), aliquoted into prechilled sterile cryovials, and frozen at −80 °C in Mr. Frosty containers (Nalgene) filled with isopropanol before transfer to liquid nitrogen for long-term storage.

### Nuclear lysis

Cryopreserved samples were rapidly thawed in a 37 °C water bath and transferred to 15 mL Falcon tubes containing 14 mL of prewarmed RPMI + 10% FBS (Thermo Fisher Scientific). After centrifugation at 350 g for 8 min at room temperature, the supernatant was removed and cell pellets were washed with 10 mL of ice-cold 1× PBS (Thermo Fisher Scientific) supplemented with 0.05% bovine serum albumin (BSA; PN 130-091-376, Miltenyi Biotec).

Cells were passed through a 40 µm cell strainer (PN 43-10040-70) and counted and assessed for viability using a TC20™ automated cell counter (Bio-Rad). Dead cells and ambient DNA were removed by flow cytometry, gating on DAPI⁻ cells on a FACSAria™ Fusion sorter (BD Biosciences).

Nuclei isolation was performed following the 10x Genomics protocol “Nuclei Isolation for Single Cell Multiome ATAC + Gene Expression” (CG000365). In brief, viable cells were pelleted by centrifugation and resuspended in 100 µL of 0.1× lysis buffer, incubating for 3 min. Nuclei were washed three times in 1× nuclei buffer supplemented with RNase inhibitor to a final concentration of 1 U/µL (PN 3335402001, Roche). After filtration through a 40 µm Flowmi strainer (PN BAH136800040-50EA, Merck), nuclei were stained with trypan blue and manually counted using a Neubauer chamber.

### Library preparation and sequencing

Gene expression (GEX) and chromatin accessibility (ATAC) libraries were prepared using the Chromium Next GEM Single Cell Multiome ATAC + Gene Expression kit according to the manufacturer’s user guide (10x Genomics, CG000338). Nuclei were subjected to transposition with the kit transposase mix, which cuts DNA at accessible chromatin regions and incorporates adapters at fragment ends. After transposition, nuclei were encapsulated into gel bead-in-emulsion (GEMs) on a Chromium Controller using a chip J, targeting ∼7,000 recovered nuclei per sample. During GEM generation, each nucleus was labeled with a 16 bp barcode and each mRNA molecule received a 12 bp unique molecular identifier (UMI) during cDNA synthesis.

GEMs were incubated to perform reverse transcription of mRNA and labeling of transposed DNA fragments. Barcoded cDNA and genomic DNA were purified and preamplified by 7 PCR cycles according to the 10x Genomics protocol. Subsequently, PCR products were purified and 35 µL of preamplified cDNA were further amplified in 7 additional PCR cycles. cDNA was quantified using a High Sensitivity DNA chip on a Bioanalyzer (Agilent Technologies), and 100 ng were used to generate GEX libraries, which were indexed with 13 PCR cycles using the Dual Index Plate TT Set A (10x Genomics; PN-3000431).

In parallel, 40 µL of preamplified genomic DNA were used to prepare ATAC libraries, indexed in 8 PCR cycles with the Sample Index N Set A (10x Genomics; PN-3000427). The size distribution and concentration of final GEX and ATAC libraries were assessed using the High Sensitivity DNA chip on a Bioanalyzer.

GEX libraries were sequenced on a NovaSeq 6000 platform (Illumina) using a 28 bp (read 1) + 10 bp (i7 index) + 10 bp (i5 index) + 90 bp (read 2) configuration, yielding >20,000 paired-end reads per cell. ATAC libraries were sequenced on a NovaSeq 6000 using 50 bp (read 1N) + 8 bp (i7 index) + 16 bp (i5 index) + 49 bp (read 2N), targeting >25,000 reads per nucleus.

Nuclear lysis and preparation of single-cell multiomic libraries from the murine spleen, followed by sequencing, were performed at the National Center for Genomic Analysis (CNAG) at the Barcelona Science Park.

### Single-cell multiome data processing

FASTQ raw files of gene expression and chromatin accessibility data from each individual library were aligned to the mouse reference genome mm10 (version mm10-2020-A-2.0.0) using Cell Ranger Arc v2.0.0 software from 10x Genomics with default parameters. A total of 81,768 filtered barcodes obtained from Cell Ranger were used for downstream analysis. The Seurat (v5.0.2)^130^ and Signac (v1.13.0)^131^ R packages were employed to process snRNA-Seq and snATAC-Seq data, respectively.

Each sample was preprocessed independently. Low-quality cells were filtered out based on the following criteria: cells with fewer than 500 UMIs, 300 expressed genes, or 500 accessible fragments were removed. Additionally, cells with excessive reads (over 20,000 UMIs, 6,000 genes, or 100,000 fragments) were excluded to prevent the inclusion of homotypic doublets or multiplets. Cells were further excluded if they exhibited more than 20% mitochondrial gene content, over 10% of fragments mapping to blacklist regions defined by ENCODE, a nucleosome signal above 2, TSS enrichment below 2, or less than 20% of regions overlapping peaks. Heterotypic doublets were detected and filtered separately for each modality using scDblFinder (v1.16.0). After applying these filters, 58,512 high-quality cells remained with an average of 3,193 UMIs and 10,073 ATAC fragments per cell (Figure S6).

### Processing of gene expression data (scRNA-seq)

scRNA-seq preprocessing was performed per sample using Seurat. Genes expressed in fewer than 10 cells, as well as genes associated with ribosomal contamination and low quality (*Malat1*, *Gm42418*, *Gm26017* and *Ay036118*), were removed. Ambient mRNA contamination was estimated and corrected with SoupX (v1.6.2) using autoEstCont() and adjustCounts() functions, and corrected matrices were used for downstream analyses. Gene expression data were first normalized using NormalizeData() function with the default LogNormalize method, and the 2,000 most highly variable genes (HVGs) per sample were selected using FindVariableFeatures() function with vst method. Cell cycle phase was estimated with CellCycleScoring(), and S and G2/M phase scores were regressed out with ScaleData(). Principal component analysis (PCA) was performed using RunPCA() on the selected HVGs, and the top 50 principal components (PCs) were retained for further analysis. Data integration across individual samples was conducted using Harmony (v1.2.3)^132^. A low-dimensional embedding was generated using the Uniform Manifold Approximation and Projection (UMAP) algorithm, applied to the first 50 integrated PCs.

### Processing of chromatin accessibility data (scATAC-seq)

Low-quality cells were filtered out before performing peak calling on each sample library with MACS3 (v3.0.0b1) via the CallPeaks() function from the Signac package, specifying an effective genome size of 1.87 × 10**⁹** bp for mouse. Peaks located outside standard chromosomes or overlapping blacklisted regions were removed. Peaks from all samples were merged into a common peak set using reduce() from GenomicRanges (v1.54.1), and accessibility matrices were recomputed with FeatureMatrix() using the common peak set and Cell Ranger fragment files as input.

scATAC-seq data were normalized per sample by latent semantic indexing (LSI). Initially, ATAC matrices were normalized using Term Frequency-Inverse Document Frequency (TF-IDF). Dimensionality reduction was then performed on the TF-IDF matrix using Singular Value Decomposition (SVD) to obtain LSI components. For visualization, a UMAP embedding was generated using the first 50 reduced dimensions, excluding the first one because it mainly reflected library size).

To integrate ATAC samples, individual objects were merged, and TF-IDF normalization and LSI reduction were repeated. Integration anchors across samples were identified with FindIntegrationAnchors() function (reciprocal LSI, dimensions 2-50), integrated embeddings were obtained with IntegrateEmbeddings(), and a final UMAP was computed from the integrated LSI dimensions.

### Integration of multimodal data

After processing snRNA-Seq and snATAC-Seq data independently, we performed weighted nearest neighbor (WNN) integration using Seurat to identify the nearest neighbors across both modalities. The WNN graph was computed using the FindMultimodalNeighbors function using the 1:50 dimensions from snRNA-Seq after harmony integration and the 2:50 integrated LSI reduction dimensions from snATAC-Seq. A joint UMAP based on this wsnn graph was then computed and used for visualization and clustering.

### Cell clustering and manual cell-type annotation

Cell-type identification was performed using a hierarchical annotation strategy. At each level, cells were processed as described for scRNA-seq and scATAC-seq (selection of variable genes/peaks, normalization, dimensionality reduction, sample integration, and multimodal WNN integration).

Clustering was performed on the wsnn graph using FindClusters() with the Smart Local Moving (SLM) algorithm, adjusting the resolution parameter according to the level and cell type to obtain biologically coherent clusters. Low-quality clusters (low FRiP or technical signal) and clusters with incompatible marker expression patterns, consistent with heterotypic doublets, were removed.

Cluster marker genes were identified with FindAllMarkers() (Wilcoxon rank-sum test), considering genes expressed in ≥10% of cells in at least one cluster and declaring significance for genes with log₂FC > 0 and FDR < 0.05. Mitochondrial and ribosomal genes were excluded from the differential expressed genes (DEGs). At the first level, 11 major clusters were identified, and one was discarded for low quality. At subsequent levels, cells were subdivided into B cells, T/NK cells, and myeloid cells; the T/NK subset was further split into CD4⁺ T cells, CD8⁺ T cells, and NK cells. In total, 48 cellular subpopulations were defined. Manual annotations were validated by automatic labeling using SingleR (v2.4.1), using as reference the murine ImmGen immune cell atlas.

### Identification of accessible regions

To capture subpopulation-specific accessibility patterns, peak calling was repeated on pseudobulk profiles generated from each cell cluster. Fragments from all cells belonging to a given cluster were aggregated, and CallPeaks() was run with Signac as in the global analysis to call cluster-specific peaks. Peak sets from all clusters were merged to obtain a final catalog of 156,396 accessible regions used for downstream analyses.

### Differentially accessible regions between cell types

Differentially accessible regions between cell types were identified using a logistic regression (LR) model implemented in FindAllMarkers() function including the number of accessible fragments per cell as a covariate. Only peaks present in ≥10% of cells in at least one cluster were considered. Peaks with log₂FC > 0 and FDR < 0.05 were defined as differentially accessible regions (DARs).

### Analysis of transcription factor activity and footprinting

TF activity was inferred from scATAC-seq data using chromVAR (v1.24.0)^32^. Motif matrices were obtained from the JASPAR2022 (v0.99.7)^133^ vertebrate core collection with getMatrixSet() from TFBSTools (v1.40.0) together with the BSgenome.Mmusculus.UCSC.mm10 (v1.4.3) package used to load the mouse genome sequences. Motifs were mapped to accessibility peaks with AddMotifs() in Signac. RunChromVAR() was then used to compute bias-corrected Z deviation scores for each motif in each cell. To identify TFs with subpopulation-specific activity, the wilcoxauc() function from presto (v1.0.0) was applied both to TF gene expression and to chromVAR Z scores, generating area under the curve (AUC) values used to rank TFs characteristic of each cluster.

TF occupancy profiles (footprints) were computed with Signac. After correcting for Tn5 insertion bias with InsertionBias(), observed and expected insertion frequencies around TF binding sites (TFBS) were calculated and footprint profiles were generated with Footprint() using only motifs located in accessible peaks. Footprint profiles were visualized per subpopulation with PlotFootprint(), allowing comparison of relative TF occupancy across cell types.

### cis-regulatory interactions between accessible regions and target genes

cis interactions between accessible regions (peaks) and putative target genes were inferred with LinkPeaks() in Signac. Spearman correlations were computed between peak accessibility and the expression of genes located within ±500 kb of the transcription start site (TSS). For each peak, a null distribution of expected correlations was generated by sampling 200 peaks with similar properties (GC content, accessibility, and length) located on other chromosomes. Peak-gene pairs with positive correlation (R > 0) and p < 0.05 were retained as candidate cis-regulatory element (CRE)-gene links.

### Identification of chromatin co-accessibility domains

Chromatin co-accessibility between genomic regions was analyzed with Cicero (v1.3.9)^36^. The scATAC-seq Seurat object was converted to a CellDataSet using as.cell_data_set() from SeuratWrappers (v0.3.4). The run_cicero() function estimated co-accessibility between peaks using a graphical LASSO model, with a maximum genomic distance of 500 kb.

To mitigate scATAC-seq sparsity, Cicero aggregates cells with similar profiles into pseudobulks of ∼50 cells via a k-nearest neighbors (KNN) strategy. Based on the co-accessibility matrix, we identified cis co-accessible chromatin neighborhoods (CCANs) using generate_ccans(), considering peak pairs with co-accessibility >0.2. CCANs were called both in the full dataset and separately within each experimental condition.

### Module scores for gene expression and chromatin accessibility

Module scores for predefined gene and peak sets were calculated with AddModuleScore() function from Seurat. We evaluated modules of canonical marker genes and peaks for each cell type, as well as signatures of germinal center precursor B cells described by King *et al.*^38^ and gene sets associated with extrafollicular, and germinal center memory B cells described by Viant *et al.*^17^ and Callahan *et al*.,^45^.

### Differential gene expression and accessibility between experimental conditions

Changes in gene expression and chromatin accessibility between primary infection and reinfection were analyzed with FindMarkers() within each cellular subpopulation. Comparisons with <50 cells in either group were excluded. Only genes or peaks detected in ≥10% of cells in at least one group were tested. For scRNA-seq, differential expression was assessed with the Wilcoxon rank-sum test. Genes with FDR < 0.05 and |log₂FC| > 0.5 were defined as DEGs, excluding mitochondrial and ribosomal genes. For scATAC-seq, a logistic regression model correcting for the number of accessible fragments per cell was used; peaks with FDR < 0.20 were considered DARs.

Differential TF activity was also evaluated with FindMarkers() using chromVAR Z scores as input, applying the same filters and specifying mean.fxn = rowMeans and fc.name = "avg_diff"; TFs with FDR < 0.20 were deemed significant.

### Functional enrichment analyses associated with gene expression changes

Functional enrichment of DEGs was assessed by over-representation analysis (ORA) using clusterProfiler (v4.10.1), with Gene Ontology (GO) Biological Process terms and KEGG pathways retrieved from org.Mm.eg.db (v. 3.18.0). Upregulated genes in primary infection or reinfection were analyzed separately within each subpopulation, using all detected genes as background. Terms containing ≥3 genes and with FDR < 0.05 were considered significantly enriched.

To reduce redundancy in GO results, we used rrvgo (v1.14.2), which groups terms according to semantic similarity (threshold 0.9) and selects the term with the lowest FDR as representative for each group. In addition, a gene set enrichment analysis (GSEA) with clusterProfiler was performed in memory B-cell subpopulations using gene sets associated with extrafollicular and germinal center differentiation described by Viant *et al.*^17^ and Callahan *et al.*^45^.

### Trajectory analysis

Trajectory analyses in B-cell subpopulations were performed with monocle3 (v1.3.7)^46^. Cycling B cells and plasmablasts were removed prior to trajectory construction. Seurat objects were converted to CellDataSet (cds) objects and UMAP coordinates derived from integrated scATAC-seq were used as the embedding space.

Trajectories were fitted with learn_graph() (use_partition = FALSE) to obtain a single continuous graph. Pseudotime was assigned with order_cells(), setting progenitor B cells (ProB) as the root state. Trajectories were reconstructed independently for each experimental condition.

### Trajectory-associated changes in expression and accessibility

Trajectory-associated changes in gene expression and accessibility were analyzed with tradeSeq (v1.16.0)^134^. Pseudotime values were rescaled to the [0,1] interval across trajectories. For gene expression, a generalized additive model was fitted with fitGAM(), including genes with >5 counts in ≥10 cells, excluding mitochondrial and ribosomal genes and using six knots. associationTest() was applied to evaluate the association of each gene with pseudotime. Genes with FDR < 0.05 and an average log₂FC along the trajectory >0.5 were defined as trajectory-associated genes (TAGs).

Differences in expression dynamics between conditions were evaluated with conditionTest() function from tradeSeq and genes with FDR < 0.01 were defined as differential trajectory-associated genes (DTAGs). An analogous analysis was performed using chromVAR Z scores to identify trajectory-associated TFs (TATFs; FDR < 0.05 and average log₂FC > 1) and TFs with differential trajectory association between conditions (DTATFs; FDR < 0.01).

### Construction of gene regulatory networks from multimodal data

Gene regulatory networks (GRNs) were constructed with GRaNIE (v1.9.7)^47^ using multimodal data aggregated into pseudobulks along B-cell trajectories. First, cells were grouped into clusters using FindClusters() on the wsnn graph (SLM algorithm, resolution = 5), and clusters with <50 cells were removed, yielding ∼50 clusters per trajectory. For each cluster, average gene expression and chromatin accessibility were computed with AverageExpression(), generating pseudobulk matrices used as GRaNIE input.

TF binding sites (TFBS) were identified within accessible peaks using addTFBS() with JASPAR2022 motifs. Spearman correlations between TF expression and accessibility of TFBS-containing peaks were then computed with addConnections_TF_peak(), correcting for GC content. GRaNIE estimated FDR thresholds in 40 correlation intervals of width 0.05 spanning −1 to 1 by comparing peaks with and without TFBS for each TF and defined significant TF-peak connections as those with FDR < 0.3.

Peak-gene links were obtained by correlating accessibility of peaks located within ±250 kb of the TSS with gene expression (Spearman), retaining peak-gene pairs with p < 0.05. The final GRN was generated by integrating significant TF-peak and peak-gene connections, linking each TF to the accessible regions it binds and to its putative target genes. TF importance within the network was quantified by PageRank centrality using page_rank() from the igraph package (v2.1.4).

## Resource Availability

### Data Availability

The single-cell multiomic data from spleens of Py17XNL-infected mice will be made publicly available in the ENA Nucleotide Archive upon publication.

## Acknowledgements

A.R.P. acknowledge funding from the Regional Programme of Research and Technological Innovation for Young Doctors UCM-CAM (PR65/19-22460). A.R.P. acknowledge funding granted by the Regional Programme of Research of “*Atracción de Talento de la Comunidad de Madrid*” (2017-T2/BMD-5532). M.C. acknowledge funding granted by the Spanish Ministry of Science, Innovation and Universities with further support from FEDER and UE (PID2022-138673OB-C21). A.V.N. acknowledge funding granted by the Regional Programme for Predoctoral Research FPI-CAM (A281-CT65/22). We thank the staff and the computing resources and technical support provided by at the SCBI (Supercomputing and Bioinformatics) Center of the University of Málaga in the Spanish Supercomputing Network (RES).

## Author Contributions

A.R.P. and JM.B. conceived the study. A.R.P., JM.B., IG.A., A.P. and A.D. designed the experiments of infection. ARP, JM.B, S.P.B. and IG.A. conceived and prepared samples for single-cell assays. M.C., A.V.N. and A.R.P. preformed the computational and bioinformatic analyses. M.C. and A.R.P. prepared the figures. A.R.P. supervised the computational analyses. A.R.P. and M.C. wrote the original draft of the manuscript with help of JM.B. All authors contributed to draft discussion, manuscript review and revisions. A.R.P., JM.B. and A.P. managed the project administration. A.R.P. and JM.B. provided fundings for the project. All authors contributed to the preparation of the paper and approved the submitted version.

---

**Figure S1.**
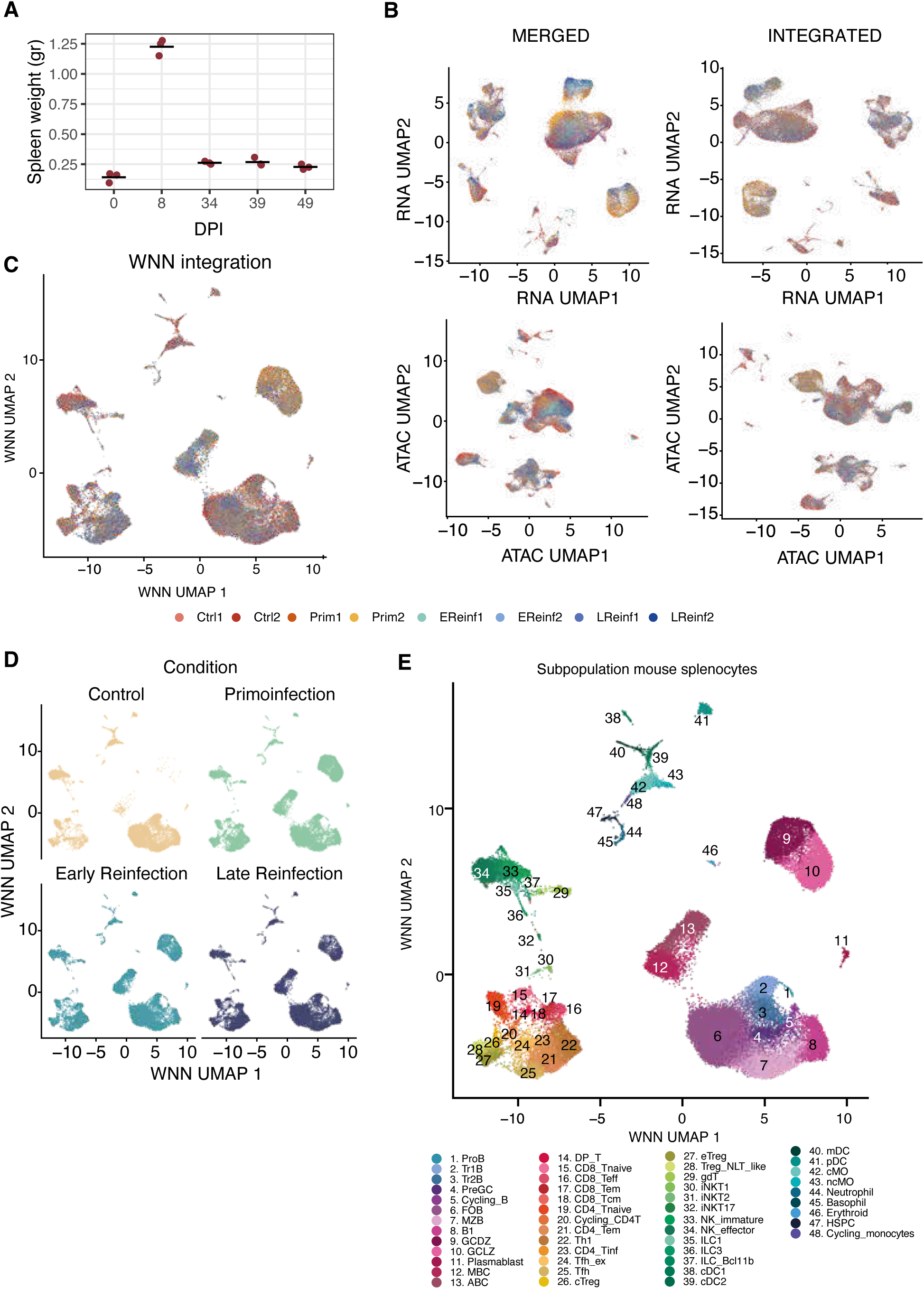
Integration of multimodal single-cell data. (A) Spleen weight measured on day 0 (uninfected group) and at 8, 34, 39 and 49 dpi. (B) UMAP representation obtained after concatenation (left) and subsequent integration (right) of scRNA-seq (top) and scATAC-seq (bottom) data from the different libraries. (C) UMAP representation after multimodal integration of both omics using WNN. Each dot represents a single cell, and color indicates the corresponding sample or condition. (D) WNN-based multimodal UMAPs split by infection group. (E) UMAP representation of the WNN-integrated dataset colored by subpopulation annotation level (n = 48 subpopulations).

**Figure S2.**
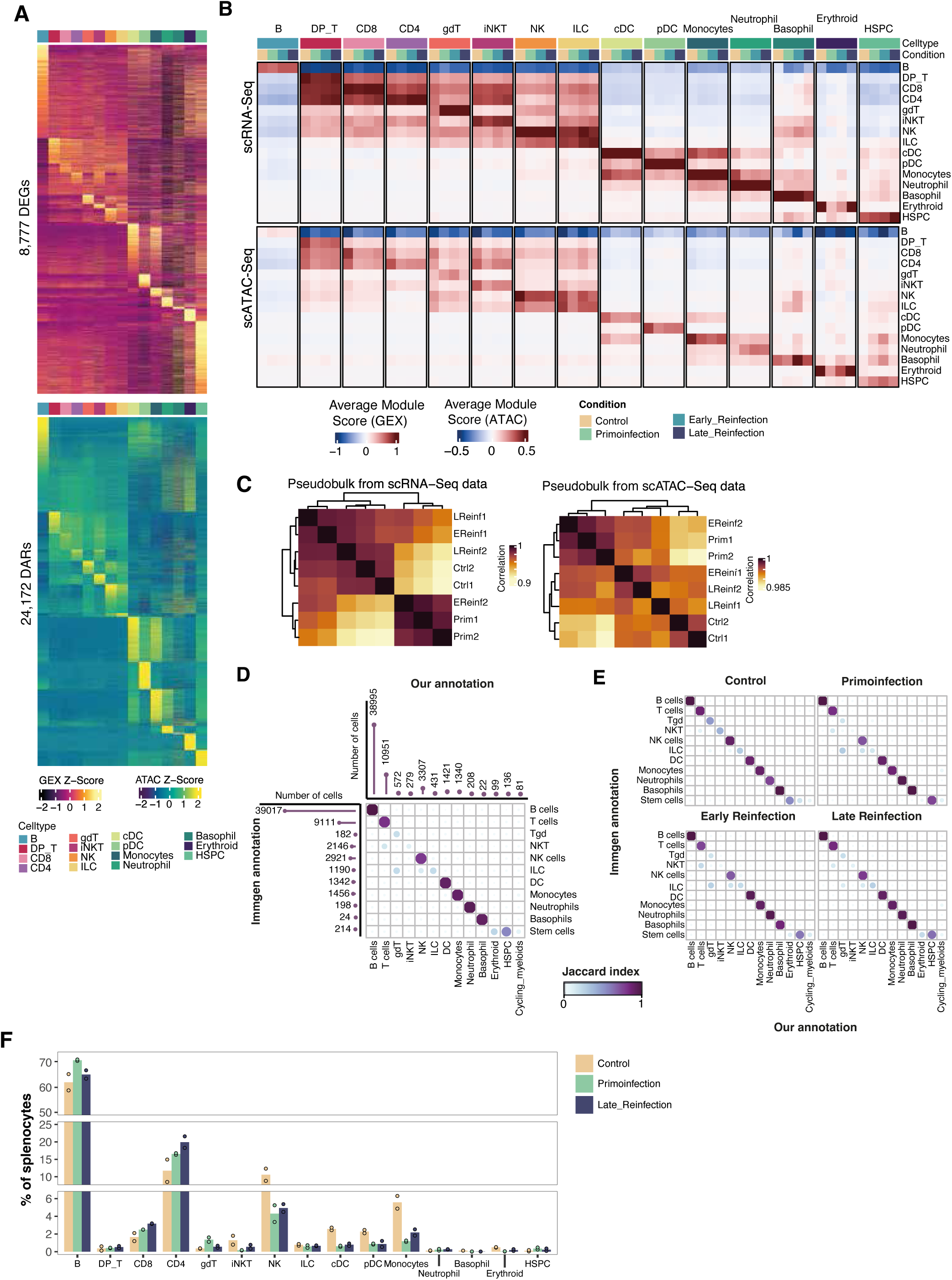
Cell-type characterization and condition-associated changes in the murine spleen across Py17XNL infection states. (A) Gene expression and accessibility profiles in each cell type. Heatmaps showing gene expression (top) and chromatin accessibility (bottom) levels for population-specific markers. Each column corresponds to a pseudobulk profile generated for a given cell type, and each row represents a gene or an accessibility peak. The color gradient indicates standardized expression and accessibility values. (B) Heatmap showing the mean module scores of marker sets derived from scRNA-seq (top) and scATAC-seq (bottom), stratified by cell type and infection group. The color gradient represents the mean module score in each combination. (C) Hierarchical clustering of scRNA-seq libraries (left) and scATAC-seq libraries (right) based on Pearson correlation. Color encodes Pearson correlation coefficients for each pairwise comparison. (D-E) Validation of major cell-type annotations. Similarity between our manual cell-type annotation and the automatic annotation obtained using the ImmGen reference database. Circle size and color indicate the Jaccard index for each comparison using all cells (D) and after stratification by experimental condition. (E) The lollipop plot in panel A indicates the number of cells annotated in each cell type. (F) Barplot showing the percentage of each cell population in individual mice, grouped by experimental condition. Each bar represents the mean percentage for the two mice within the same infection group, while circles indicate individual values.

**Figure S3.**
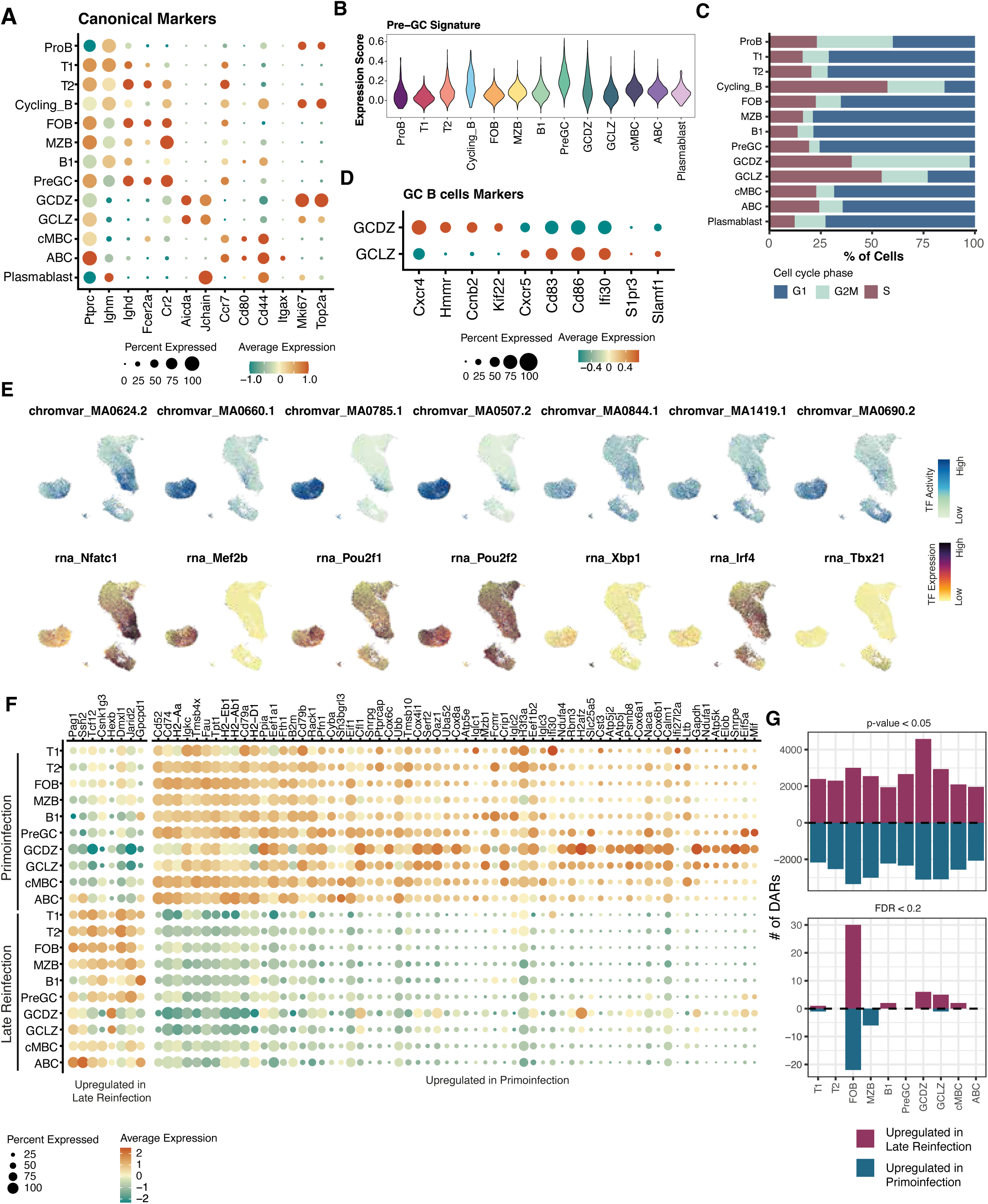
Single-cell multiomic features of B cell subpopulation. (A) Mean expression of B cell canonical marker genes. (B) Violin plots showing the distribution of normalized expression values of set of PreGC gene^38^ in individual cells, grouped by B cell subpopulation. (C) Percentage of cells in each cell-cycle phase (G1, G2/M, and S) for each B cell subpopulation. (D) Mean expression of marker genes used to define germinal center B cells^40^. (E) Characterization of TF activity in B lymphocytes. UMAP plots showing TF activity (top) and expression (bottom) for TFs specific to the different B cell subpopulations. (F) Expression of the 71 DEGs common to all B cell subpopulations, shown by subpopulation and infection group. The color gradient reflects normalized, standardized gene expression, and dot size indicates the proportion of cells expressing each gene in each subpopulation. (G) Chromatin accessibility changes associated with the number of Py17XNL exposures in B cell subpopulations. Barplots showing the number of DARs with increased (log2FC > 0) or decreased (log2FC < 0) accessibility in the comparison of reinfection versus primary infection, separated into regions with nominal p-value < 0.05 (left) and those with FDR < 0.2 (right).

**Figure S4.**
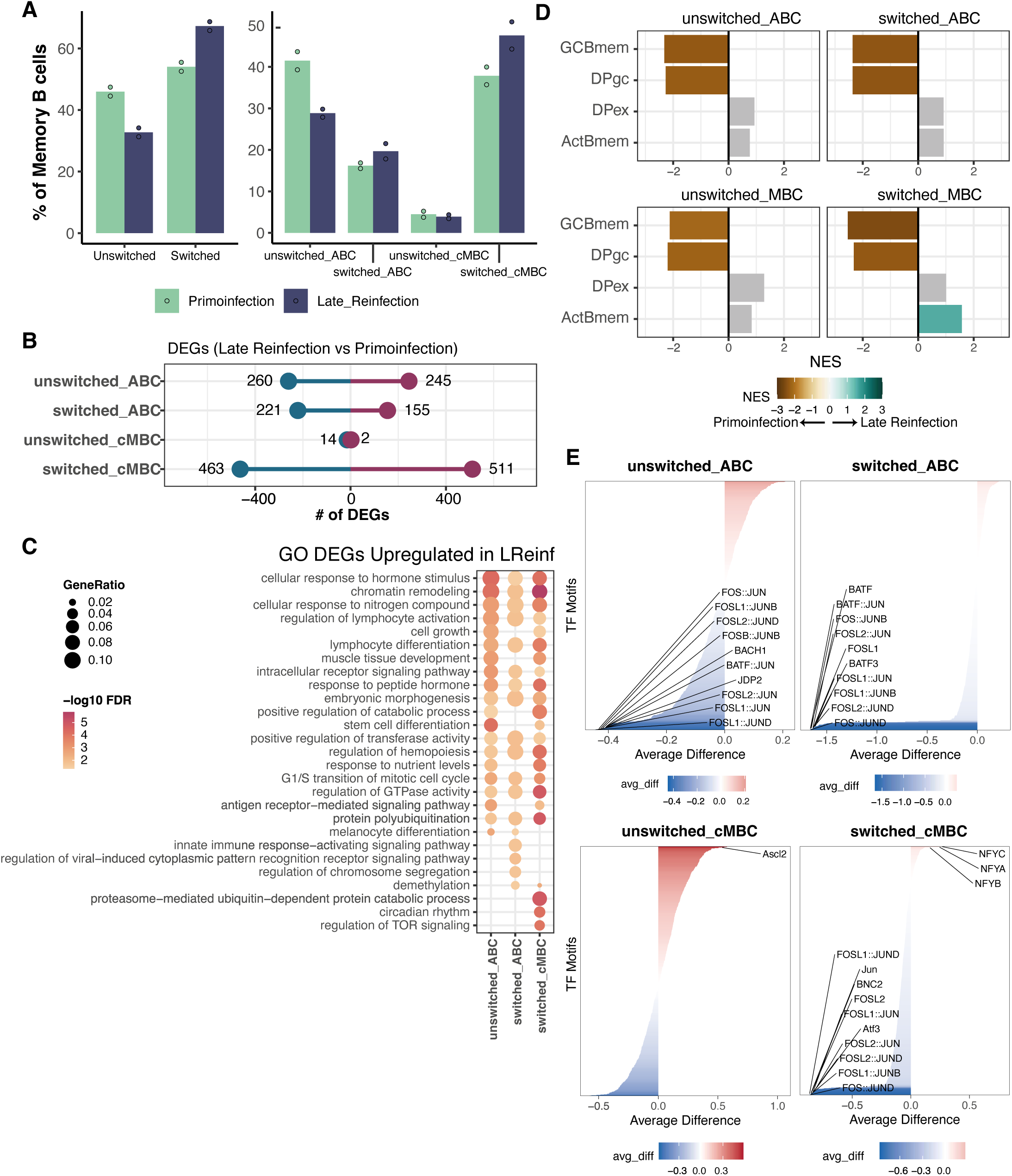
Heterogeneity of memory B cells after multiple exposures to Py17XNL. (A) Percentage of each memory B cell subpopulation in individual mice, grouped by experimental condition. Bars indicate the mean percentage for each infection group, and circles show individual mouse values. (B) Number of DEGs identified in each memory B cell subpopulation in the comparison of reinfection versus primary infection. Red circles denote DEGs overexpressed in reinfection, and blue circles denote DEGs overexpressed in primary infection. (C) Top 10 enriched GO biological process terms based on DEGs overexpressed in reinfection compared to primary infection; circle size is proportional to the ratio between DEGs and the total number of genes in each enriched term, and the color gradient indicates −log10(FDR). Circles highlight the memory B cell subpopulation in which each term is enriched. (D) Barplots showing GSEA results using gene signatures for extrafollicular- and GC-derived memory B cells from Viant *et al*.^17^ and Callahan *et al.*^45^, computed from log2FC values for reinfection versus primary infection in each memory B cell subpopulation. The x-axis and color gradient represent the normalized enrichment score (NES). Gene sets with NES > 0 (green) are enriched in reinfection; gene sets with NES < 0 (brown) are enriched in primary infection. Grey bars indicate non-significant gene sets (p-value > 0.05). (E) Changes in TF activity when comparing reinfection versus primary infection in each memory B cell subpopulation. The 10 TFs with the highest activity in each condition are shown. The color gradient indicates higher activity in primary infection (blue) or reinfection (red).

**Figure S5.**
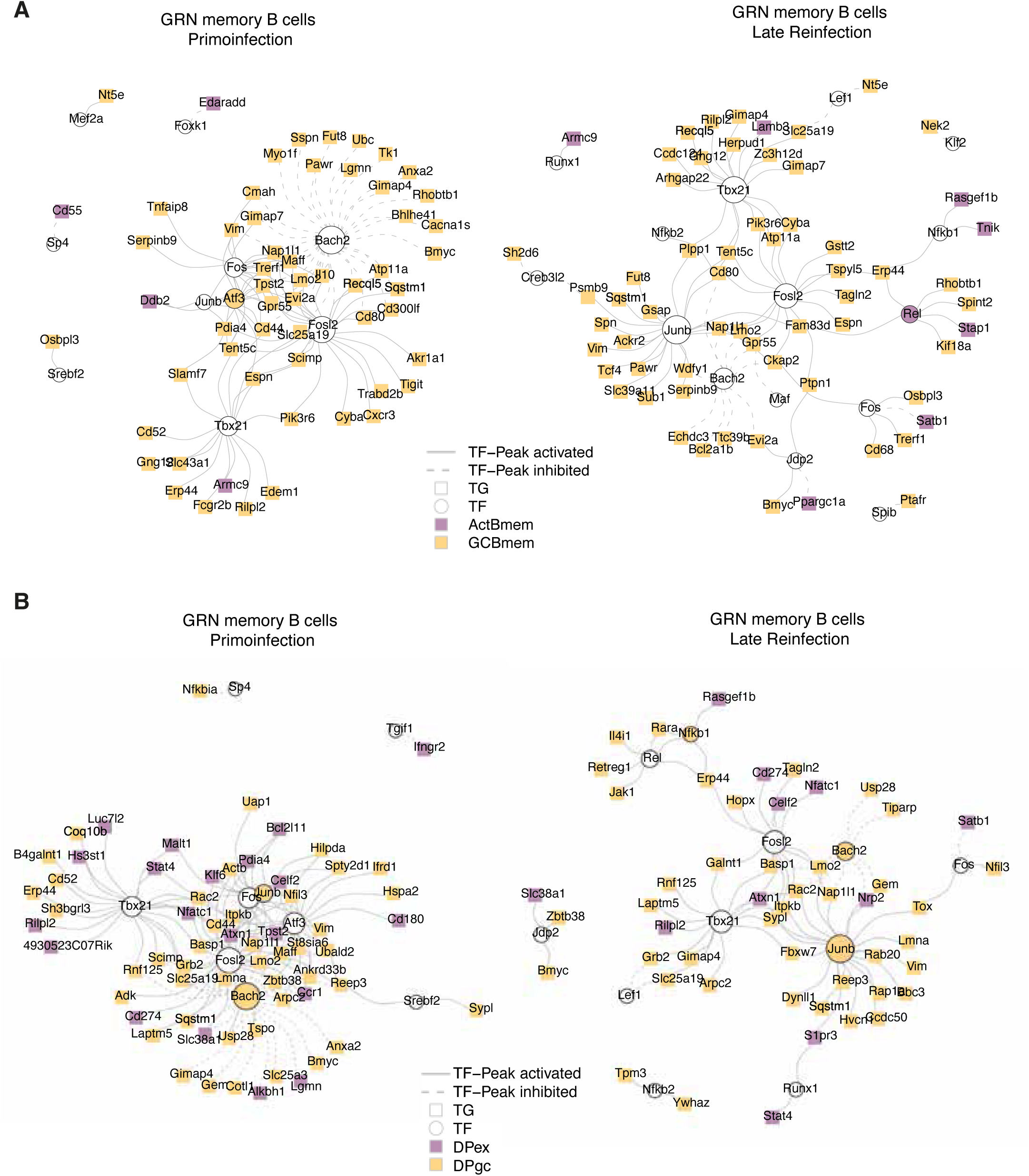
Identification of genes associated with memory B cell differentiation in gene regulatory networks. GRNs of target genes associated with differentiation of memory B cells derived from extrafollicular or GC origin, using gene sets from Viant *et al.*^17^ (A) and Callahan *et al.* ^45^ (B), shown for primary infection GRNs (left) and reinfection GRNs (right). Activating interactions are shown as solid lines and repressive interactions as dashed lines. Circles represent TFs and squares represent target genes. Genes associated with extrafollicular differentiation are shown in purple, and genes associated with GC-derived differentiation are shown in yellow.

**Figure S6.**
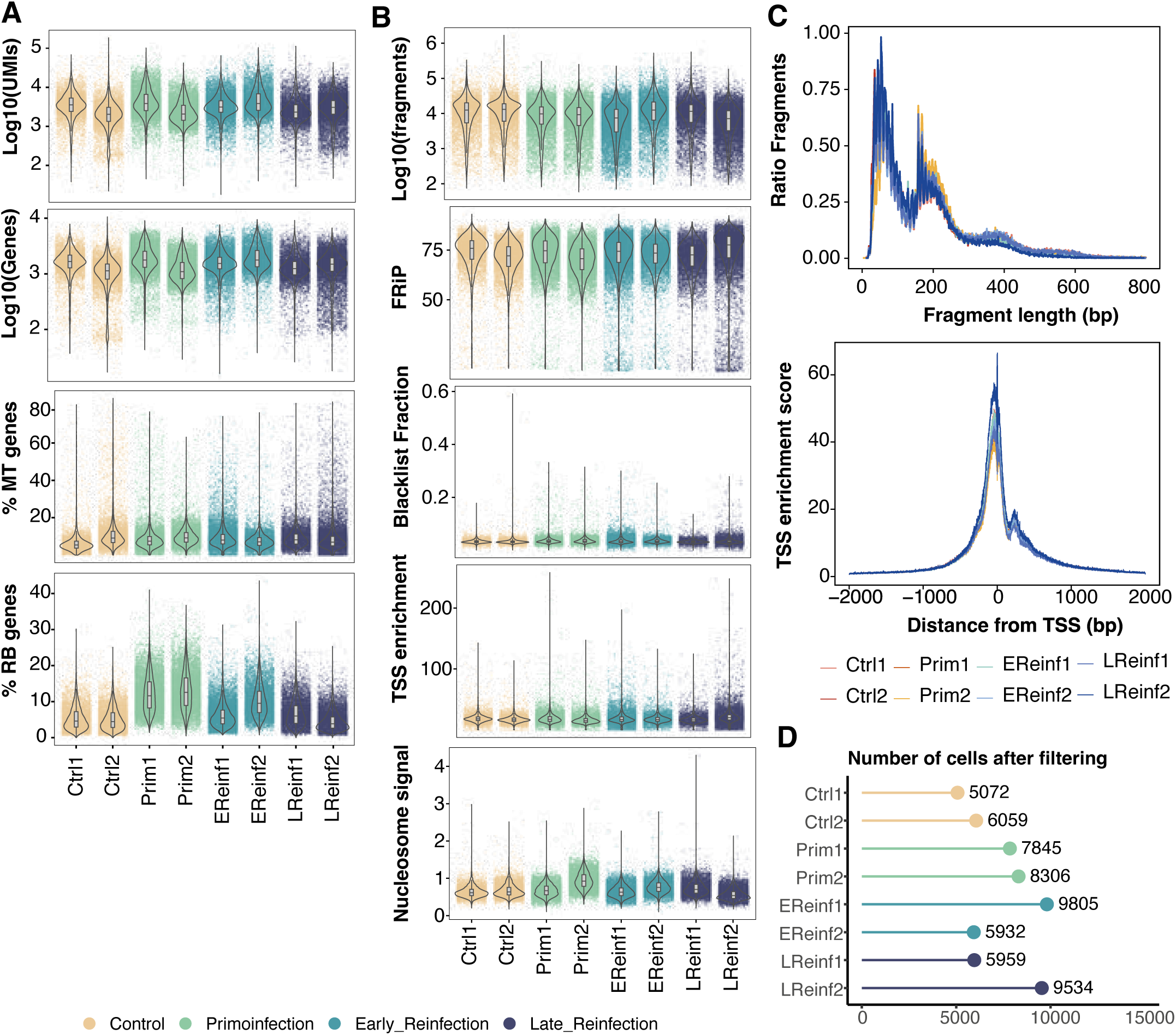
Quality control of scRNA-Seq and scATAC-seq data per sample. (A) Boxplots showing RNA-seq quality metrics per cell, including the number of reads (UMIs), the number of expressed genes, and the percentage of mitochondrial and ribosomal reads, separated by sample and colored by the infected group. (B) Boxplots displaying ATAC-seq quality metrics, including fragment count, fraction of fragments in peaks (FRiP), fraction of peaks in blacklisted regions, TSS enrichment, and nucleosome signal. (C) Distributions of fragment sizes for each scATAC-seq sample (top) and aggregated normalized fragment counts around TSSs (bottom) for each scATAC-seq sample. (D) Number of cells per sample after filtering out low-quality cells.

**Table S1.** Top 15 upregulated DEGs across B-Cell subpopulations.

| Cluster | Genes |
| --- | --- |
| **ProB** | Sox4, 2610307P16Rik, Gfra1, Tnfrsf19, Epb41l4b, Vpreb1, Adgrg1, Erg, Gfra2, Il7r, Atp1b1, Cacna2d1, Cplx2, 2900026A02Rik, B230118H07Rik |
| **Tr1B** | Agbl1, Mctp2, Ighm, Ppm1e, Serinc5, Eps8, Map2, Aff3, Foxp1, Tmem163, Cecr2, Ptprj, Cux1, Sox4, Fam129c |
| **Tr2B** | Lef1, 2010309G21Rik, Cdk6, Agbl1, Foxn3, Foxp1, Ssbp2, Arhgap24, Pard3b, Aff3, Pip5k1b, Il12a, Ighd, Myb, Dmxl1 |
| **Cycling_B** | Cntln, Alpl, Slc43a3, Rad51b, Grn, Brip1, Cenpp, Diaph3, Cit, Cpd, Gm42047, Polq, Ncapg2, Pard3b, Bard1 |
| **FOB** | Satb1, Ighd, Lrrk2, Cd55, Zfp318, Fcer2a, Gpr183, Tnik, Rasgef1b, Xylt1, Akt3, Stat4, Abca1, Cpne4, Notch2 |
| **MZB** | Atxn1, Pik3r4, Cr2, Plaur, Trps1, St6galnac3, Maml2, Camkmt, Mndal, Zdhhc14, Ptprj, Lrrk2, Rasgef1b, Zc3h12c, Utrn |
| **B1** | Cacna1e, Atxn1, Il5ra, Agbl1, 1810046K07Rik, Plac8, Ptpn22, Bcl2, Pstpip2, Il6st, Nfatc1, Tbc1d9, Ubash3b, Ahnak, Rbm47 |
| **PreGC** | Mdn1, Hsp90ab1, Ccnd2, Ncl, Arhgap24, Rel, Gnl3, March3, Pus7, Hspd1, Mybbp1a, Npm1, Mettl1, Hspa5, Atad3a |
| **GCDZ** | Mki67, Top2a, Kif11, Pclaf, Diaph3, Ect2, Rgs13, Basp1, Cenpe, Stmn1, Mybl1, Tox, Ezh2, Tuba1b, Knl1 |
| **GCLZ** | Rgs13, Tox, Pdzd2, Mybl1, Basp1, Mtf2, Jchain, Slc41a2, Parp8, Rfc1, Neil1, Hmces, Trerf1, Aicda, Pitpnc1 |
| **MBc** | Ahnak, Parm1, Cmah, Sspn, Cd44, Vim, Syne1, Gimap4, Myo1f, Phtf2, Cd80, Tnfrsf8, Gimap7, Gm32312, Gpr183 |
| **ABC** | Zeb2, Adgre1, Pld4, Ahnak, Cd44, Cd72, Vps37b, Arhgef18, Ptpn22, Fcgr2b, Itgb1, Dclre1c, Heg1, Plaur, Gns |
| **Plasmablast** | Sec24d, Derl3, Rhobtb1, Creb3l2, Prdm1, Cpeb3, Ly6c2, Ckap4, Iglv1, Dnm3, Sdc1, Fkbp11, Prlr, Ly6c1, Hid1 |

## References

1. Ryg-Cornejo, V., Ly, A. & Hansen, D. S. Immunological processes underlying the slow acquisition of humoral immunity to malaria. Parasitology 143, 199–207 (2016).

2. Langhorne, J., Ndungu, F. M., Sponaas, A.-M. & Marsh, K. Immunity to malaria: more questions than answers. Nat Immunol 9, 725–732 (2008).

3. Weiss, G. E. et al. The Plasmodium falciparum-Specific Human Memory B Cell Compartment Expands Gradually with Repeated Malaria Infections. PLoS Pathog 6, e1000912 (2010).

4. Ferrer, P. et al. Repeat controlled human Plasmodium falciparum infections delay bloodstream patency and reduce symptoms. Nat Commun 15, 5194 (2024).

5. Tran, T. M. et al. An Intensive Longitudinal Cohort Study of Malian Children and Adults Reveals No Evidence of Acquired Immunity to Plasmodium falciparum Infection. Clinical Infectious Diseases 57, 40–47 (2013).

6. Ghosh, D. & Stumhofer, J. S. The spleen: “epicenter” in malaria infection and immunity. Journal of Leukocyte Biology 110, 753–769 (2021).

7. Henry, B. et al. The Human Spleen in Malaria: Filter or Shelter? Trends in Parasitology 36, 435–446 (2020).

8. Henry, B. et al. Splenic clearance of rigid erythrocytes as an inherited mechanism for splenomegaly and natural resistance to malaria. eBioMedicine 82, 104167 (2022).

9. Huang, X. et al. Differential Spleen Remodeling Associated with Different Levels of Parasite Virulence Controls Disease Outcome in Malaria Parasite Infections. mSphere 1, e00018–15 (2016).

10. Lewis, S. M., Williams, A. & Eisenbarth, S. C. Structure and function of the immune system in the spleen. Sci. Immunol. 4, eaau6085 (2019).

11. Alves, F. A. et al. Splenic architecture disruption and parasite-induced splenocyte activation and anergy in Plasmodium falciparum-infected Saimiri sciureus monkeys. Malar J 14, 128 (2015).

12. Wang, Q. et al. Adaptive immune responses mediated age-related Plasmodium yoelii 17XL and 17XNL infections in 4 and 8-week-old BALB/c mice. BMC Immunol 22, 6 (2021).

13. Chauhan, R. et al. CD4+ICOS+Foxp3+: a sub-population of regulatory T cells contribute to malaria pathogenesis. Malar J 21, 32 (2022).

14. Lee, H. J. et al. CD4+ T cells display a spectrum of recall dynamics during re-infection with malaria parasites. Nat Commun 15, 5497 (2024).

15. Calôba, C. et al. Systemic 4-1BB stimulation augments extrafollicular memory B cell formation and recall responses during Plasmodium infection. Cell Reports 44, 115528 (2025).

16. Kurosaki, T., Kometani, K. & Ise, W. Memory B cells. Nat Rev Immunol 15, 149–159 (2015).

17. Viant, C. et al. Germinal center–dependent and –independent memory B cells produced throughout the immune response. Journal of Experimental Medicine 218, e20202489 (2021).

18. Valeri, V. et al. B cell intrinsic and extrinsic factors impacting memory recall responses to SRBC challenge. Front. Immunol. 13, 873886 (2022).

19. Budeus, B., Kibler, A. & Küppers, R. Human IgM–expressing memory B cells. Front. Immunol. 14, 1308378 (2023).

20. Syeda, M. Z., Hong, T., Huang, C., Huang, W. & Mu, Q. B cell memory: from generation to reactivation: a multipronged defense wall against pathogens. Cell Death Discov. 10, 117 (2024).

21. Portugal, S. et al. Malaria-associated atypical memory B cells exhibit markedly reduced B cell receptor signaling and effector function. eLife 4, e07218 (2015).

22. Pérez-Mazliah, D. et al. Plasmodium-specific atypical memory B cells are short-lived activated B cells. eLife 7, e39800 (2018).

23. Mouat, I. C., Goldberg, E. & Horwitz, M. S. Age-associated B cells in autoimmune diseases. Cell. Mol. Life Sci. 79, 402 (2022).

24. Ricker, E. et al. Altered function and differentiation of age-associated B cells contribute to the female bias in lupus mice. Nat Commun 12, 4813 (2021).

25. Krishnamurty, A. T. et al. Somatically Hypermutated Plasmodium-Specific IgM+ Memory B Cells Are Rapid, Plastic, Early Responders upon Malaria Rechallenge. Immunity 45, 402–414 (2016).

26. Brown, S. L., Bauer, J. J., Lee, J., Ntirandekura, E. & Stumhofer, J. S. IgM+ and IgM– memory B cells represent heterogeneous populations capable of producing class-switched antibodies and germinal center B cells upon rechallenge with *P. yoelii*. Journal of Leukocyte Biology 112, 1115–1135 (2022).

27. Dooley, N. L. et al. Single cell transcriptomics shows that malaria promotes unique regulatory responses across multiple immune cell subsets. Nat Commun 14, 7387 (2023).

28. Sundling, C. et al. B cell profiling in malaria reveals expansion and remodeling of CD11c+ B cell subsets. JCI Insight 4, e126492 (2019).

29. Zheng, G. X. Y. et al. Massively parallel digital transcriptional profiling of single cells. Nat Commun 8, 14049 (2017).

30. Satpathy, A. T. et al. Massively parallel single-cell chromatin landscapes of human immune cell development and intratumoral T cell exhaustion. Nat Biotechnol 37, 925–936 (2019).

31. Hao, Y. et al. Integrated analysis of multimodal single-cell data. Cell 184, 3573–3587.e29 (2021).

32. Schep, A. N., Wu, B., Buenrostro, J. D. & Greenleaf, W. J. chromVAR: inferring transcription-factor-associated accessibility from single-cell epigenomic data. Nat Methods 14, 975–978 (2017).

33. The Immunological Genome Project Consortium et al. The Immunological Genome Project: networks of gene expression in immune cells. Nat Immunol 9, 1091–1094 (2008).

34. Ma, S. et al. Chromatin Potential Identified by Shared Single-Cell Profiling of RNA and Chromatin. Cell 183, 1103–1116.e20 (2020).

35. Bour-Jordan, H. et al. Intrinsic and extrinsic control of peripheral T-cell tolerance by costimulatory molecules of the CD28/ B7 family. Immunological Reviews 241, 180–205 (2011).

36. Pliner, H. A. et al. Cicero Predicts cis-Regulatory DNA Interactions from Single-Cell Chromatin Accessibility Data. Molecular Cell 71, 858–871.e8 (2018).

37. Sun, B. et al. Sox4 Is Required for the Survival of Pro-B Cells. The Journal of Immunology 190, 2080–2089 (2013).

38. King, H. W., et al. Single-cell analysis of human B cell maturation predicts how antibody class switching shapes selection dynamics. Sci. Immunol. 6, eabe6291 (2021).

39. Mathew, N. R. et al. Single-cell BCR and transcriptome analysis after influenza infection reveals spatiotemporal dynamics of antigen-specific B cells. Cell Reports 35, 109286 (2021).

40. Victora, G. D. et al. Germinal Center Dynamics Revealed by Multiphoton Microscopy with a Photoactivatable Fluorescent Reporter. Cell 143, 592–605 (2010).

41. Gao, X., et al. Zeb2 drives the formation of CD11c^+^ atypical B cells to sustain germinal centers that control persistent infection. Sci. Immunol. 9, eadj4748 (2024).

42. Li, G. et al. Jarid2 and PRC2, partners in regulating gene expression. Genes Dev. 24, 368–380 (2010).

43. Kinkel, S. A. et al. Jarid2 regulates hematopoietic stem cell function by acting with polycomb repressive complex 2. Blood 125, 1890–1900 (2015).

44. Al-Raawi, D. et al. A novel form of JARID2 is required for differentiation in lineage-committed cells. The EMBO Journal 38, e98449 (2019).

45. Callahan, D. et al. Memory B cell subsets have divergent developmental origins that are coupled to distinct imprinted epigenetic states. Nat Immunol 25, 562–575 (2024).

46. Cao, J. et al. The single-cell transcriptional landscape of mammalian organogenesis. Nature 566, 496–502 (2019).

47. Kamal, A. et al. GRaNIE and GRaNPA: inference and evaluation of enhancer-mediated gene regulatory networks. Molecular Systems Biology 19, e11627 (2023).

48. Langhorne, J. The immune response to the blood stages of Plasmodium in animal models. Immunology Letters 41, 99–102 (1994).

49. Kalkal, M. & Das, J. Differential B cell mediated immune response during Plasmodium yoelii infection in mice. Acta Tropica 263, 107533 (2025).

50. Fu, Y., Ding, Y., Zhou, T., Fu, X. & Xu, W. Plasmodium yoelii blood-stage primes macrophage-mediated innate immune response through modulation of toll-like receptor signalling. Malar J 11, 104 (2012).

51. Imai, T. et al. Involvement of CD8^+^ T cells in protective immunity against murine blood-stage infection with *Plasmodium yoelii* 17XL strain. Eur J Immunol 40, 1053–1061 (2010).

52. Muxel, S. M. et al. The Spleen CD4+ T Cell Response to Blood-Stage Plasmodium chabaudi Malaria Develops in Two Phases Characterized by Different Properties. PLoS ONE 6, e22434 (2011).

53. Stephens, R., Culleton, R. L. & Lamb, T. J. The contribution of Plasmodium chabaudi to our understanding of malaria. Trends in Parasitology 28, 73–82 (2012).

54. Achtman, A. H., Stephens, R., Cadman, E. T., Harrison, V. & Langhorne, J. Malaria-specific antibody responses and parasite persistence after infection of mice with *Plasmodium chabaudi chabaudi*. Parasite Immunology 29, 435–444 (2007).

55. Ma, S. et al. Plasmodium yoelii: Influence of antimalarial treatment on acquisition of immunity in BALB/c and DBA/2 mice. Experimental Parasitology 116, 266–272 (2007).

56. Imai, T. et al. Involvement of CD8^+^ T cells in protective immunity against murine blood-stage infection with *Plasmodium yoelii* 17XL strain. Eur J Immunol 40, 1053–1061 (2010).

57. Oster, C. N., Koontz, L. C. & Wyler, D. J. Malaria in Asplenic Mice: Effects of Splenectomy, Congenital Asplenia, and Splenic Reconstitution on the Course of Infection. The American Journal of Tropical Medicine and Hygiene 29, 1138–1142 (1980).

58. Otto, M. I., Vliegenthart-Jongbloed, K. J., Van Hellemond, J. J. & Van Genderen, P. J. Plasmodium falciparum malaria runs a more severe course in splenectomized patients at comparable levels of parasitemia: a retrospective matched case-control study. Trop Dis Travel Med Vaccines 11, 18 (2025).

59. Bachmann, A. et al. Absence of Erythrocyte Sequestration and Lack of Multicopy Gene Family Expression in Plasmodium falciparum from a Splenectomized Malaria Patient. PLoS ONE 4, e7459 (2009).

60. Carvalho, L. J., Ferreira-da-Cruz, M. F., Daniel-Ribeiro, C. T., Pelajo-Machado, M. & Lenzi, H. L. Germinal center architecture disturbance during Plasmodium berghei ANKA infection in CBA mice. Malar J 6, 59 (2007).

61. Achtman, A. H., Khan, M., MacLennan, I. C. M. & Langhorne, J. *Plasmodium chabaudi chabaudi* Infection in Mice Induces Strong B Cell Responses and Striking But Temporary Changes in Splenic Cell Distribution. The Journal of Immunology 171, 317–324 (2003).

62. Chen, S. et al. The Dynamic Change of Immune Responses Between Acute and Recurrence Stages of Rodent Malaria Infection. Front. Microbiol. 13, 844975 (2022).

63. Williams, C. G. et al. Plasmodium infection induces phenotypic, clonal, and spatial diversity among differentiating CD4+ T cells. Cell Reports 43, 114317 (2024).

64. Skinner, O. P. et al. Temporally overlapping mechanisms diversify clonal B cell responses *in vivo*. Preprint at 10.1101/2024.12.17.628863 (2024).

65. Hensel, J. A., Khattar, V., Ashton, R. & Ponnazhagan, S. Characterization of immune cell subtypes in three commonly used mouse strains reveals gender and strain-specific variations. Laboratory Investigation 99, 93–106 (2019).

66. Hoetzel, K., Feuerstein, H., Ludwig, J. & Wardemann, H. Multi-parameter spectral flow cytometry panel for immune phenotyping of murine B and T cell responses. Preprint at 10.1101/2025.07.18.665522 (2025).

67. Hansen, D. S. et al. CD1d-restricted NKT cells contribute to malarial splenomegaly and enhance parasite-specific antibody responses. Eur J Immunol 33, 2588–2598 (2003).

68. Azcárate, I. G. et al. Early and late B cell immune responses in lethal and self-cured rodent malaria. Immunobiology 220, 684–691 (2015).

69. Nduati, E. W. et al. Distinct Kinetics of Memory B-Cell and Plasma-Cell Responses in Peripheral Blood Following a Blood-Stage Plasmodium chabaudi Infection in Mice. PLoS ONE 5, e15007 (2010).

70. Ndungu, F. M. et al. Functional Memory B Cells and Long-Lived Plasma Cells Are Generated after a Single Plasmodium chabaudi Infection in Mice. PLoS Pathog 5, e1000690 (2009).

71. Anderson, S. M., Tomayko, M. M. & Shlomchik, M. J. Intrinsic properties of human and murine memory B cells. Immunological Reviews 211, 280–294 (2006).

72. Sutton, H. J. et al. Atypical B cells are part of an alternative lineage of B cells that participates in responses to vaccination and infection in humans. Cell Reports 34, 108684 (2021).

73. Moir, S. et al. Evidence for HIV-associated B cell exhaustion in a dysfunctional memory B cell compartment in HIV-infected viremic individuals. The Journal of Experimental Medicine 205, 1797–1805 (2008).

74. Wei, C. et al. A New Population of Cells Lacking Expression of CD27 Represents a Notable Component of the B Cell Memory Compartment in Systemic Lupus Erythematosus. The Journal of Immunology 178, 6624–6633 (2007).

75. Muellenbeck, M. F. et al. Atypical and classical memory B cells produce *Plasmodium falciparum* neutralizing antibodies. Journal of Experimental Medicine 210, 389–399 (2013).

76. Kim, C. C., Baccarella, A. M., Bayat, A., Pepper, M. & Fontana, M. F. FCRL5+ Memory B Cells Exhibit Robust Recall Responses. Cell Reports 27, 1446–1460.e4 (2019).

77. Gao, X., et al. Zeb2 drives the formation of CD11c^+^ atypical B cells to sustain germinal centers that control persistent infection. Sci. Immunol. 9, eadj4748 (2024).

78. Arnold, C. N. et al. A forward genetic screen reveals roles for *Nfkbid*, *Zeb1*, and *Ruvbl2* in humoral immunity. Proc. Natl. Acad. Sci. U.S.A. 109, 12286–12293 (2012).

79. Papadopoulou, V., Postigo, A., Sánchez-Tilló, E., Porter, A. C. G. & Wagner, S. D. ZEB1 and CtBP form a repressive complex at a distal promoter element of the BCL6 locus. Biochemical Journal 427, 541–550 (2010).

80. Vijay, R. et al. Infection-induced plasmablasts are a nutrient sink that impairs humoral immunity to malaria. Nat Immunol 21, 790–801 (2020).

81. Stephens, R., Ndungu, F. M. & Langhorne, J. Germinal centre and marginal zone B cells expand quickly in a second *Plasmodium chabaudi* malaria infection producing mature plasma cells. Parasite Immunology 31, 20–31 (2009).

82. Huse, K. et al. Mechanism of CD79A and CD79B Support for IgM+ B Cell Fitness through B Cell Receptor Surface Expression. The Journal of Immunology 209, 2042–2053 (2022).

83. Tolar, P. Cytoskeletal control of B cell responses to antigens. Nat Rev Immunol 17, 621–634 (2017).

84. Vardhana, S. A. et al. Impaired mitochondrial oxidative phosphorylation limits the self-renewal of T cells exposed to persistent antigen. Nat Immunol 21, 1022–1033 (2020).

85. Yao, C.-H. et al. Mitochondrial fusion supports increased oxidative phosphorylation during cell proliferation. eLife 8, e41351 (2019).

86. Pasini, D. et al. JARID2 regulates binding of the Polycomb repressive complex 2 to target genes in ES cells. Nature 464, 306–310 (2010).

87. Toyoda, M. et al. jumonji Downregulates Cardiac Cell Proliferation by Repressing cyclin D1 Expression. Developmental Cell 5, 85–97 (2003).

88. Shirato, H. et al. A Jumonji (Jarid2) Protein Complex Represses cyclin D1 Expression by Methylation of Histone H3-K9. Journal of Biological Chemistry 284, 733–739 (2009).

89. Cao, R. et al. Role of Histone H3 Lysine 27 Methylation in Polycomb-Group Silencing. Science 298, 1039–1043 (2002).

90. Müller, J. et al. Histone Methyltransferase Activity of a Drosophila Polycomb Group Repressor Complex. Cell 111, 197–208 (2002).

91. Simon, M. et al. Single-cell chromatin accessibility and transposable element landscapes reveal shared features of tissue-residing immune cells. Immunity 57, 1975–1993.e10 (2024).

92. Schwartz, Y. B. et al. Genome-wide analysis of Polycomb targets in Drosophila melanogaster. Nat Genet 38, 700–705 (2006).

93. Simon, J. A. & Kingston, R. E. Occupying Chromatin: Polycomb Mechanisms for Getting to Genomic Targets, Stopping Transcriptional Traffic, and Staying Put. Molecular Cell 49, 808–824 (2013).

94. Raaphorst, F. M. et al. Cutting Edge: Polycomb Gene Expression Patterns Reflect Distinct B Cell Differentiation Stages in Human Germinal Centers. The Journal of Immunology 164, 1–4 (2000).

95. Van Galen, J. C. et al. Distinct expression patterns of polycomb oncoproteins and their binding partners during the germinal center reaction. Eur J Immunol 34, 1870–1881 (2004).

96. Dogan, I. et al. Multiple layers of B cell memory with different effector functions. Nat Immunol 10, 1292–1299 (2009).

97. Pape, K. A., Catron, D. M., Itano, A. A. & Jenkins, M. K. The Humoral Immune Response Is Initiated in Lymph Nodes by B Cells that Acquire Soluble Antigen Directly in the Follicles. Immunity 26, 491–502 (2007).

98. Ambegaonkar, A. A. et al. Isotype switching in human memory B cells sets intrinsic antigen-affinity thresholds that dictate antigen-driven fates. Proc. Natl. Acad. Sci. U.S.A. 121, e2313672121 (2024).

99. Seifert, M. et al. Functional capacities of human IgM memory B cells in early inflammatory responses and secondary germinal center reactions. Proc. Natl. Acad. Sci. U.S.A. 112, (2015).

100. Seifert, M. et al. Functional capacities of human IgM memory B cells in early inflammatory responses and secondary germinal center reactions. Proc. Natl. Acad. Sci. U.S.A. 112, (2015).

101. LeBien, T. W. & Tedder, T. F. B lymphocytes: how they develop and function. Blood 112, 1570–1580 (2008).

102. Yosef, N. & Regev, A. Writ large: Genomic dissection of the effect of cellular environment on immune response. Science 354, 64–68 (2016).

103. Busslinger, M. & Tarakhovsky, A. Epigenetic Control of Immunity. Cold Spring Harbor Perspectives in Biology 6, a019307–a019307 (2014).

104. Giltiay, N. V., Giordano, D. & Clark, E. A. The Plasticity of Newly Formed B Cells. The Journal of Immunology 203, 3095–3104 (2019).

105. Wei, C., Yu, P. & Cheng, L. Hematopoietic Reprogramming Entangles with Hematopoiesis. Trends in Cell Biology 30, 752–763 (2020).

106. Chen, S., Yang, J., Wei, Y. & Wei, X. Epigenetic regulation of macrophages: from homeostasis maintenance to host defense. Cell Mol Immunol 17, 36–49 (2020).

107. Trapnell, C. Defining cell types and states with single-cell genomics. Genome Res. 25, 1491–1498 (2015).

108. Cannoodt, R., Saelens, W. & Saeys, Y. Computational methods for trajectory inference from single-cell transcriptomics. Eur J Immunol 46, 2496–2506 (2016).

109. Liebermann, D. A., Gregory, B. & Hoffman, B. AP-1 (Fos/Jun) transcription factors in hematopoietic differentiation and apoptosis. Int J Oncol 10.3892/ijo.12.3.685 (1998) doi:10.3892/ijo.12.3.685.

110. Grötsch, B. et al. The AP-1 transcription factor Fra1 inhibits follicular B cell differentiation into plasma cells. Journal of Experimental Medicine 211, 2199–2212 (2014).

111. Ma, S. et al. Chromatin Potential Identified by Shared Single-Cell Profiling of RNA and Chromatin. Cell 183, 1103–1116.e20 (2020).

112. Fleck, J. S. et al. Inferring and perturbing cell fate regulomes in human brain organoids. Nature 621, 365–372 (2023).

113. Bravo González-Blas, C., et al. SCENIC+: single-cell multiomic inference of enhancers and gene regulatory networks. Nat Methods 20, 1355–1367 (2023).

114. Kartha, V. K. et al. Functional inference of gene regulation using single-cell multi-omics. Cell Genomics 2, 100166 (2022).

115. Reyes-Palomares, A. et al. Systematic identification of phenotypically enriched loci using a patient network of genomic disorders. BMC Genomics 17, 232 (2016).

116. Daga, N. et al. Integration of genetic and chromatin modification data pinpoints autoimmune-specific remodeling of enhancer landscape in CD4+ T cells. Cell Rep 43, 114810 (2024).

117. Lobato-Moreno, S. et al. Single-cell ultra-high-throughput multiplexed chromatin and RNA profiling reveals gene regulatory dynamics. Nat Methods 22, 1213–1225 (2025).

118. Kometani, K. et al. Repression of the Transcription Factor Bach2 Contributes to Predisposition of IgG1 Memory B Cells toward Plasma Cell Differentiation. Immunity 39, 136–147 (2013).

119. Hai, T. The ATF Transcription Factors in Cellular Adaptive Responses. in Gene Expression and Regulation (ed. Ma, J.) 329–340 (Springer New York, New York, NY, 2006). doi:10.1007/978-0-387-40049-5_20.

120. Ku, H.-C. & Cheng, C.-F. Master Regulator Activating Transcription Factor 3 (ATF3) in Metabolic Homeostasis and Cancer. Front. Endocrinol. 11, 556 (2020).

121. Yang, S.-Y. et al. Characterization of Organ-Specific Regulatory B Cells Using Single-Cell RNA Sequencing. Front. Immunol. 12, 711980 (2021).

122. Gilchrist, M. et al. Systems biology approaches identify ATF3 as a negative regulator of Toll-like receptor 4. Nature 441, 173–178 (2006).

123. Oeckinghaus, A. & Ghosh, S. The NF- B Family of Transcription Factors and Its Regulation. Cold Spring Harbor Perspectives in Biology 1, a000034–a000034 (2009).

124. De Silva, N. S. et al. Transcription factors of the alternative NF-κB pathway are required for germinal center B-cell development. Proc. Natl. Acad. Sci. U.S.A. 113, 9063–9068 (2016).

125. Heise, N. et al. Germinal center B cell maintenance and differentiation are controlled by distinct NF-κB transcription factor subunits. Journal of Experimental Medicine 211, 2103–2118 (2014).

126. Pohl, T. et al. The combined absence of NF-κB1 and c-Rel reveals that overlapping roles for these transcription factors in the B cell lineage are restricted to the activation and function of mature cells. Proc. Natl. Acad. Sci. U.S.A. 99, 4514–4519 (2002).

127. Shokhirev, M. N. et al. A multi-scale approach reveals that NF-κB CR el enforces a B-cell decision to divide. Molecular Systems Biology 11, 783 (2015).

128. Tumang, J. R. et al. c-Rel is essential for B lymphocyte survival and cell cycle progression. Eur. J. Immunol. 28, 4299–4312 (1998).

129. Shih, V. F.-S., Tsui, R., Caldwell, A. & Hoffmann, A. A single NFκB system for both canonical and non-canonical signaling. Cell Res 21, 86–102 (2011).

130. Hao, Y. et al. Dictionary learning for integrative, multimodal and scalable single-cell analysis. Nat Biotechnol 42, 293–304 (2024).

131. Stuart, T., Srivastava, A., Madad, S., Lareau, C. A. & Satija, R. Single-cell chromatin state analysis with Signac. Nat Methods 18, 1333–1341 (2021).

132. Korsunsky, I. et al. Fast, sensitive and accurate integration of single-cell data with Harmony. Nat Methods 16, 1289–1296 (2019).

133. Castro-Mondragon, J. A. et al. JASPAR 2022: the 9th release of the open-access database of transcription factor binding profiles. Nucleic Acids Research 50, D165–D173 (2022).

134. Van Den Berge, K., et al. Trajectory-based differential expression analysis for single-cell sequencing data. Nat Commun 11, 1201 (2020).

